# Bayesian Tip-dated Phylogenetics: Topological Effects, Stratigraphic Fit and the Early Evolution of Mammals

**DOI:** 10.1101/533885

**Authors:** Benedict King, Robin Beck

## Abstract

The incorporation of stratigraphic data into phylogenetic analysis has a long history of debate, but is not currently standard practice for palaeontologists. Bayesian tip-dating (or morphological clock) phylogenetic methods have returned these arguments to the spotlight, but how tip-dating affects the recovery of evolutionary relationships has yet to be fully explored. Here we show, through analysis of several datasets with multiple phylogenetic methods, that topologies produced by tip-dating are outliers when compared to topologies produced by parsimony and undated Bayesian methods, which retrieve broadly similar trees. Unsurprisingly, trees recovered by tip-dating have better fit to stratigraphy than trees recovered by other methods, due to trees with better stratigraphic fit being assigned a higher prior probability. Differences in stratigraphic fit and tree topology between tip-dating and other methods appear to be concentrated in parts of the tree with weaker character signal and a stronger influence of the prior, as shown by successive deletion of the most incomplete taxa from a sauropod dataset. Tip-dating applied to Mesozoic mammals firmly rejects a monophyletic Allotheria, and strongly supports diphyly of haramiyidans, with the late Triassic *Haramiyavia* and *Thomasia* forming a clade with tritylodontids, which is distant from the middle Jurassic euharamiyidans. This result is not sensitive to the controversial age of the eutherian *Juramaia*. A Test of the age of *Juramaia* using a less restrictive prior reveals strong support from the data for an Early Cretaceous age. Our results suggest that tip-dating incorporates stratigraphic data in an intuitive way, with good stratigraphic fit a prior expectation that can be overturned by strong evidence from character data.

---

The question of whether or not the stratigraphic ages of fossils should be taken into account when estimating phylogeny was a major debate within palaeontology in the 1990s and early 2000s (Wagner 1995, Lockwood 1998, Siddall 1998, Smith 1998, Fox et al. 1999, Heyning and Thacker 1999, Smith 2000, Geiger et al. 2001, Alroy 2002). A number of methods were developed to integrate stratigraphic data with phylogenetic analysis. Wagner (1995) introduced a method to find the most parsimonious cladogram in which sister taxa have overlapping 95% confidence intervals for stratigraphic range. Stratocladistics (Fisher 2008) is a parsimony method that considers stratigraphic parsimony debt alongside “standard” morphological parsimony debt. The method of Wagner (1998) finds the tree with the maximum likelihood of the observed tree length and stratigraphic debt, calculated using simulations under a defined model of evolution. None of these methods have seen widespread adoption in the palaeontological literature, perhaps partly due to a lack of user-friendly software; for example StrataPhy, a software program for stratocladistics (Marcot and Fox 2008), was never updated from its original experimental form.

Recently, the rise of Bayesian tip-dating methods (Ronquist et al. 2012a), has seen the return of phylogenetic analysis incorporating stratigraphic data. At the heart of tip-dating methods is the tree prior: the prior probability distribution of divergence dates and branch lengths. Early tip-dated analyses used the uniform tree prior (Ronquist et al. 2012a), which is relatively uninformative regarding tree shape, and can produce unrealistic results (Matzke and Wright 2016). The uniform tree prior has been superseded by serially sampled fossilised birth-death (FBD) tree priors (Stadler 2010, Heath et al. 2014), which model diversification, extinction and sampling. Recently, these have been updated to allow sampled ancestors (Gavryushkina et al. 2014). Serially sampled tree priors include assumptions of approximately constant rates of diversification, extinction and sampling, although this can be relaxed between different time slices (Stadler et al. 2013). These tree priors likely affect tree topology, and indeed assumptions of approximately constant sampling rates between lineages at specified time intervals is a key component of stratocladistics (Fisher 2008).

Tip-dating has been used on a number of datasets to infer the phylogeny of clades with fossil members, including Mesozoic birds (Lee et al. 2014b), Mesozoic mammals (Close et al. 2015), theropod dinosaurs (Lee et al. 2014a, Bapst et al. 2016), pufferfish (Close et al. 2016) and penguins (Gavryushkina et al. 2017). Most of these studies have focused on macro-evolutionary patterns and divergence dates. However, Bapst et al. (2016) reported topological differences between tip-dated Bayesian, undated Bayesian, and parsimony analyses (as well as between different implementations of tip-dating) of a theropod dataset. Turner et al. (2017) showed that tip-dating can affect tree topology in an analysis of crocodylomorphs by disfavouring long un-sampled branches, whilst King et al. (2017) found that tip-dating can have major effects on tree topology by attempting to balance inferred rates of evolution.

Stratigraphic fit measures are explicit measures for assessing how well a phylogeny fits with the order of appearance (i.e. geological ages) of its taxa. Historically, they have been put to a number of uses, including assessing the quality of the fossil record (Benton et al. 2000) and comparisons of the stratigraphic congruence between major groups (O’Connor and Wills 2016). These indices include the Stratigraphic Consistency Index (SCI), Manhattan Stratigraphic Measure (MSM) and the Gap Excess Ratio (GER). The SCI (Huelsenbeck 1994) calculates the number of consistent nodes, where a consistent node is one whose oldest descendant is the same age or younger than the oldest descendant of its sister node. The MSM (Siddall 1998) and GER (Wills 1999) both rely on measuring the Minimum Implied Gap (MIG), a measure of the smallest possible sum of ghost lineages implied by a particular tree topology.

Here, we investigate the topological differences that result when discrete morphological character matrices are analysed using parsimony and both undated and tip-dating Bayesian methods, and their relationship with stratigraphic fit, as measured by GER and SCI. We examine datasets representing a range of vertebrate and invertebrate animal groups, namely ichthyosaurs, eurypterids, horseshoe crabs, baleen whales, turtles, sauropods, Mesozoic birds, and Mesozoic mammals. We focus our discussion on the latter group, specifically on two key questions. Firstly, we test whether the different methods support monophyly of Allotheria. Putative allotherians include haramiyidans, multituberculates and gondwanatherians, all of which share a number of dental apomorphies, most notably postcanines with multiple cusps in longitudinal rows (Kielan-Jaworowska et al. 2004, Krause et al. 2014, Huttenlocker et al. 2018). Some phylogenetic analyses have supported monophyly of Allotheria, within (crown-clade) Mammalia, with haramiyidans forming a paraphyletic assemblage (Zheng et al. 2013, Bi et al. 2014, Krause et al. 2014, Chang et al. 2017, Meng et al. 2018). Conversely, others have recovered haramiyidans outside Mammalia, with multituberculates falling within Mammalia. Monophyly versus polyphyly of Allotheria has major implications for our understanding of Mesozoic mammal evolution, leading to different scenarios for the evolution of numerous dental and skeletal features, including the Definitive Mammalian Middle Ear, in which the angular, articular, prearticular, and quadrate have become entirely auditory in function, separate from the jaw joint (Meng et al. 2018). It also affects interpretations of the age of Mammalia: if late Triassic haramiyidans fall within the crown-clade, then the split between monotremes and therians must be at least this old (Bi et al. 2014); if they fall outside the crown-clade, this split could be considerably younger (depending on the affinities of other late Triassic mammaliaforms, such as *Kuehneotherium* (Luo et al. 2015).

Secondly, we use tip-dating to investigate the age of *Juramaia sinensis*, which is reportedly part of the middle Jurassic (164-159 million year old) Linglongta Biota (the younger of the two phases comprising the Yanliao/Daohugou Biota) from the Lanqi/Tiaojishan Formation (Luo et al. 2011, Xu et al. 2016). If so, *Juramaia* extends the known age range of eutherians by approximately 20-40 million years, depending on the affinities of *Durlstotherium* and *Durlstodon* from southern England (Sweetman et al. 2017). However, *Juramaia* appears remarkably “advanced” (particularly in terms of its dentition) for its reported age, closely resembling eutherians from the much younger (~126 Ma old) Yixian Formation, such as *Eomaia* and *Ambolestes* (Meng 2014, Bi et al. 2018). Several authors have raised questions about the age of *Juramaia* and it has been suggested that it instead may come from younger (Early Cretaceous) strata in China (Meng 2014, Sullivan et al. 2014, Bi et al. 2018). If it is in fact closer to 126 Ma in age, then this results in a much younger palaeontological minimum for the timing of the split between Eutheria (placentals and their fossil relatives) and Metatheria (marsupials and their fossil relatives).

## Materials and Methods

### Comparing Tree Topology and Stratigraphic Fit between Methods

We selected a number of recent datasets for testing the topological differences between parsimony, undated Bayesian (also known as non-clock) and tip-dated Bayesian methods. These were datasets of ichthyosaurs (Ji et al. 2016), eurypterids (Lamsdell and Selden 2017), horseshoe crabs (Lamsdell 2016), baleen whales (Marx and Fordyce 2015), turtles (Perea et al. 2014), sauropods (Poropat et al. 2016), Mesozoic birds (Wang et al. 2015), and Mesozoic mammals (Krause et al. 2014, Huttenlocker et al. 2018). Stratigraphic range data were taken from these publications where available, or otherwise from the Paleobiology Database (http://fossilworks.org/). Stratigraphic ranges were converted to age ranges using the International Chronostratigraphy Chart version 2016/12 (http://www.stratigraphy.org/index.php/ics-chart-timescale) and the geowhen database (http://www.stratigraphy.org/upload/bak/geowhen/index.html).

A number of taxa were pruned from the following datasets prior to analysis: for the ichthyosaurs dataset, all outgroup taxa except *Hupesuchus*; for the horseshoe crabs dataset, all non-xiphosuran taxa except some synziphosurines; for the turtles dataset, all outgroup taxa, with the tree instead rooted on *Odontochelys*; for the Mesozoic birds dataset, all modern taxa deleted; for the Mesozoic mammals dataset, all modern and fossil Cenozoic taxa. These deletions were necessary either due to uncertainties regarding outgroup taxa, or extremely poor sampling within large time divisions. Characters that became invariant following taxon pruning were deleted from the datasets, as these are uninformative in parsimony analyses and Bayesian analyses that assume that only variable characters are included (as used here).

Parsimony analyses in TNT (Goloboff et al. 2008) employed new technology search, using sectorial search and tree fusing with default settings for 1000 random addition sequences. Due to the large total number of most-parsimonious trees for some datasets, and the computational burden of downstream analyses, we saved only the set of trees output from the new technology search, without running a traditional search to find all possible most-parsimonious trees. Apomorphy lists and Templeton tests (Templeton 1983) were performed in PAUP* 4.0b10 (Swofford 2002).

Undated Bayesian analyses were performed in MrBayes (Ronquist et al. 2012b). The Mkv model (which assumes that only variable characters have been included; Lewis 2001) was used, with a gamma parameter with four rate categories to account for rate variation across sites. Four independent runs of each analysis, each with four chains, were run for 10 million generations, saving 2000 trees. Convergence of the four runs was confirmed in Tracer (Rambaut et al. 2014).

Tip-dated Bayesian analyses were performed in BEAST2 (Bouckaert et al. 2014). The Mkv model (Lewis 2001) was used, with a gamma parameter with four rate categories to account for rate variation across sites (i.e. the same implementation as in the undated MrBayes analyses). Characters were partitioned according to the number of character states, with the substitution rate reweighted following King et al. (2017). The clock model was an uncorrelated lognormal clock (Drummond et al. 2006). The tree prior was a sampled-ancestor fossilised birth-death model (Gavryushkina et al. 2014). A rho parameter was used for those datasets containing modern taxa (horseshoe crabs, whales, turtles). Rho was set to 1.0, 0.8 and 0.083 respectively based on the proportion of extant taxa included in the analysis. For the four datasets without modern taxa, sampled ancestors was turned off (by setting removal probability to 1), due to incompatibility issues with the “TipDatesRandomWalker” operator and sampled ancestors. A standard set of priors was used for all analyses, details of which are in the xml files. Analyses were run for between 200 million and 1 billion generations (depending on dataset) with 2000 trees saved. Convergence of four independent runs was checked in Tracer. Topological convergence was confirmed using RWTY (Warren et al. 2017). Each dataset was also analysed using the prior only, without character data so that the stratigraphic fit of the prior tree sample could be calculated.

To assess the extent of topological differences between the three phylogenetic methods, we calculated Robinson-Foulds distances (Robinson and Foulds 1981) in R using the package phangorn (Schliep 2010). Every tree produced by one method was compared to every tree produced from both of the other methods. Robinson-Foulds distances were rescaled to a percentage difference following Wright and Hillis (2014). In addition to comparisons between methods, all trees from each set were compared to each other as a measure of topological variation within each method.

Stratigraphic fit, using the Gap Excess Ratio (Wills 1999), was calculated using the R package strap (Bell and Lloyd 2015). The Stratigraphic Completeness Index (Huelsenbeck 1994) was also calculated, with very similar results. To avoid problems with the different treatment of outgroups between methods, outgroups were removed from all trees prior to calculation.

Because the results (see below) showed that Bayesian tip-dated methods place a higher prior probability on trees with better stratigraphic fit, we hypothesised that this prior would be particularly influential on tree topology in regions where the character data were weak. We therefore tested the effect of incomplete taxa on the stratigraphic fit of tip-dated and undated Bayesian phylogenetic trees. This was achieved through sequential removal of incomplete taxa. The sauropods dataset was chosen due to the wide range of data completeness across taxa and the large number of taxa. For each deletion iteration (total of five), we removed the six remaining most incomplete taxa and reanalysed the data reanalysed and calculated stratigraphic fit as above. As a control, to test whether or not the act of removing taxa changes stratigraphic fit regardless of the completeness of those taxa, we repeated the process but deleted six random taxa in each iteration.

### Additional Analyses on Mesozoic Mammals Datasets

To investigate conflicts between the different parts of the dataset of Huttenlocker et al. (2018), and further test allotherian relationships, the following character subsets were analysed individually, using Bayesian tip-dating approaches: craniodental, dental only, cranial only and postcranial only. To further test the extent to which a tip-dating approach could overturn topologies supported under other methods, a similar analysis was run on the Mesozoic mammals dataset of Krause et al. (2014), which shows only partial overlap in terms of characters and taxa with that of Huttenlocker et al. (2018). Notably both parsimony and undated Bayesian analysis on this dataset supported monophyly of Allotheria (= haramiyidans, gondwanatherians and multituberculates). With Bayesian tip-dating analysis of the Krause et al. (2014) dataset, very poor mixing caused precisely by the phylogenetic conflict under investigation (monophyly or polyphyly of Allotheria), meant that this analysis was run for 44 independent runs, each of a billion generations (Metropolis coupling was ineffective in improving effective sample sizes); results from each run were thinned and combined for further analysis.

We also ran an analysis of the Huttenlocker et al. (2018) dataset to test the robustness of results to uncertainties about the age of the Chinese fossil *Juramaia sinensis* (detailed above), as large uncertainty about the age of an important taxon may be detrimental to a tip-dating analysis. To test the effect (if any) of age uncertainty of *Juramaia* on the results, we re-ran the analysis with an alternative prior distribution on the age of *Juramaia*. The prior was a Laplace distribution centred on 161 Million years, but with no hard maximum or minimum bounds, thus representing a strong prior expectation that *Juramaia* is Middle Jurassic (as originally described), but allowing other dates at low prior probability.

The use of a Laplace prior with no hard bounds on the age of *Juramaia* not only tests the effect of age uncertainty on the results of the analysis, but can also been seen as a test of the age of *Juramaia*. Bayesian phylogenetic estimation of fossil ages has been shown to accurately estimate known ages in datasets for penguins and canids (Drummond and Stadler 2016). Estimates using this method can, however, be very imprecise: in a study of hominids, morphological data were not particularly informative regarding the age of *Homo naledi* (which was treated as unknown), with 95% highest posterior density intervals spanning 0 to more than 2 million years and a mean of 912 thousand years for this species (Dembo et al. 2016). Nevertheless, the true age of *H. naledi* (236 – 335 thousand years) (Dirks et al. 2017), falls within the 95% HPD calculated by Dembo et al. (2016). The similarity of *Juramaia* to eutherians from the Early Cretaceous Yixian Formation is one of the reasons doubts have been raised about its age (Meng 2014, Bi et al. 2018) and a tip-dated analysis with a wide age prior for *Juramaia* amounts to a quantitative test of this idea. As a control measure, two other taxa from the Yanliao/Daohugou Biota, *Castorocauda* and *Rugosodon*, were given the same Laplace prior.

The results of the main analysis on the Mesozoic mammals dataset suggested that haramiyidans are diphyletic, in contrast to the result from parsimony (see below). One possible explanation for this is that the tree prior places a higher prior probability on haramiyidan diphyly compared to monophyly, whereas the data does not strongly support monophyly. We tested this by running an analysis without data on a partially fixed topology (based on the main result from tip-dating) in which only haramiyidans were free to move around the tree. Specifically, a series of backbone constraints based on the maximum clade credibility tree from the main analysis were implemented, and haramiyidans were constrained to form two monophyletic groups *(Haramiyavia* + *Thomasia*, and euharamiyidans) but with their phylogenetic position otherwise unconstrained. The analysis therefore tested where these two groups attached to the backbone, based purely on stratigraphic ranges. Extraction of the relevant information in R required the packages ape (Paradis et al. 2004), phangorn (Schliep 2010) and treeio (Yu et al. 2017). Finally, a parsimony analysis with a negative constraint on haramiyidan monophyly was run in TNT to see how strongly the morphological character data supports monophyly.

## Results

### Tree Topology

Tree topology of undated Bayesian and parsimony trees were more similar to each other than either was to tip-dated Bayesian trees (Fig. 1, Fig. S1). This is further shown by plotting the number of parsimony steps for the trees produced by Bayesian methods with and without the tip-dating (Fig. 2). The output from parsimony is more resolved than the output from the Bayesian methods (fig. 1), as previously reported (O’Reilly et al. 2016). Topological differences between undated Bayesian and parsimony trees are in general no greater than the differences within the posterior sample of undated Bayesian trees. These results suggest that it is the use of tip-dating and associated tree models, rather than a model of morphological evolution, that has the most significant effect on tree topology.

**Figure 1.**
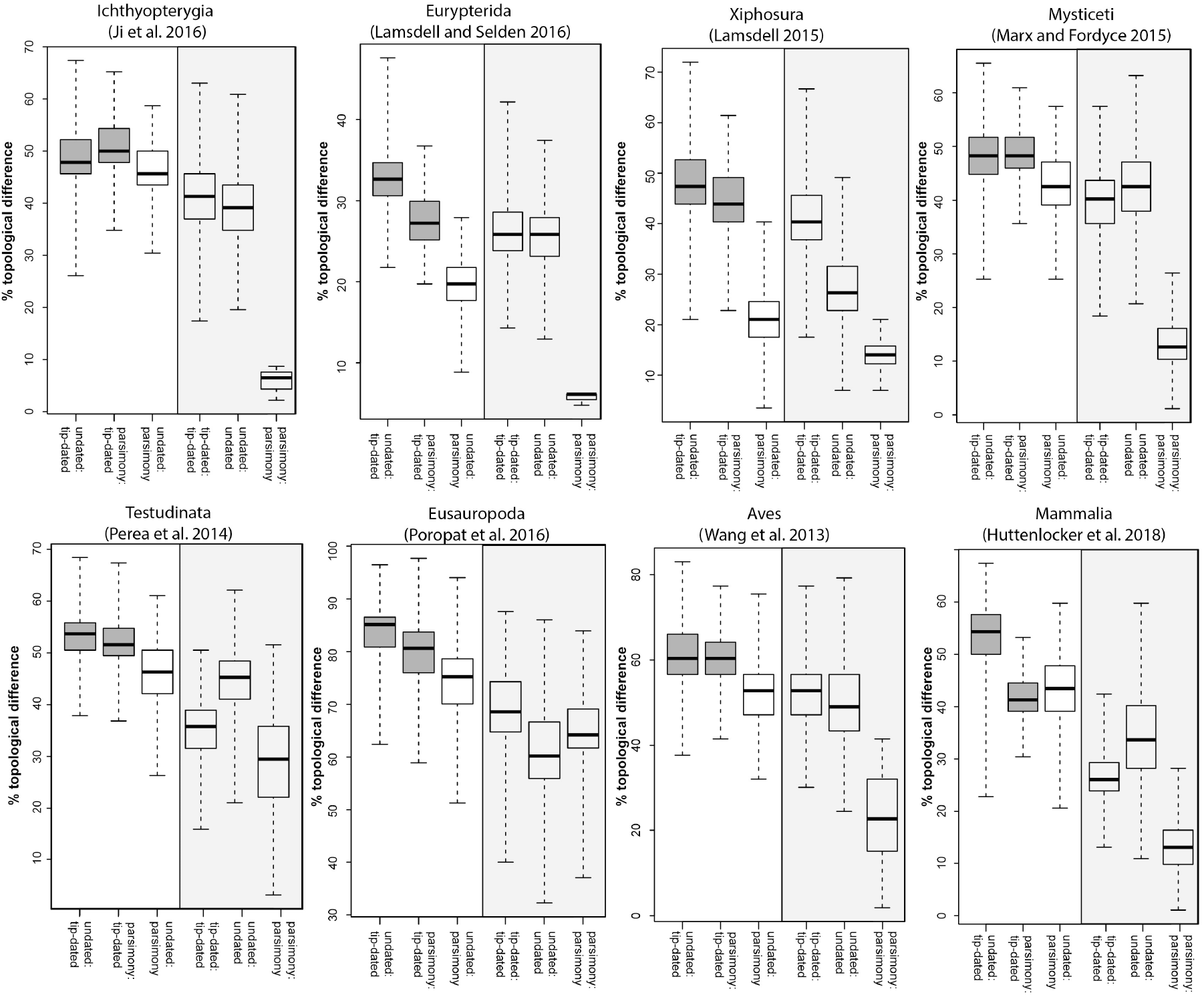
Bayesian tip-dating methods produce trees that are more different from trees produced by parsimony and undated Bayesian analyses, which are similar to each other. % topological difference (Robinson-Foulds distance) is plotted for each comparison, across seven datasets. Bayesian tip-dated methods vs. other methods (Bayesian undated and parsimony) are shaded in dark grey. Every tree from the posterior sample or set of shortest trees is compared to the sample from an alternative method, and the resulting range of values shown as a boxplot (whiskers span full range). Tree samples from each method are also compared to themselves as a measure of resolution.

**Figure 2.**
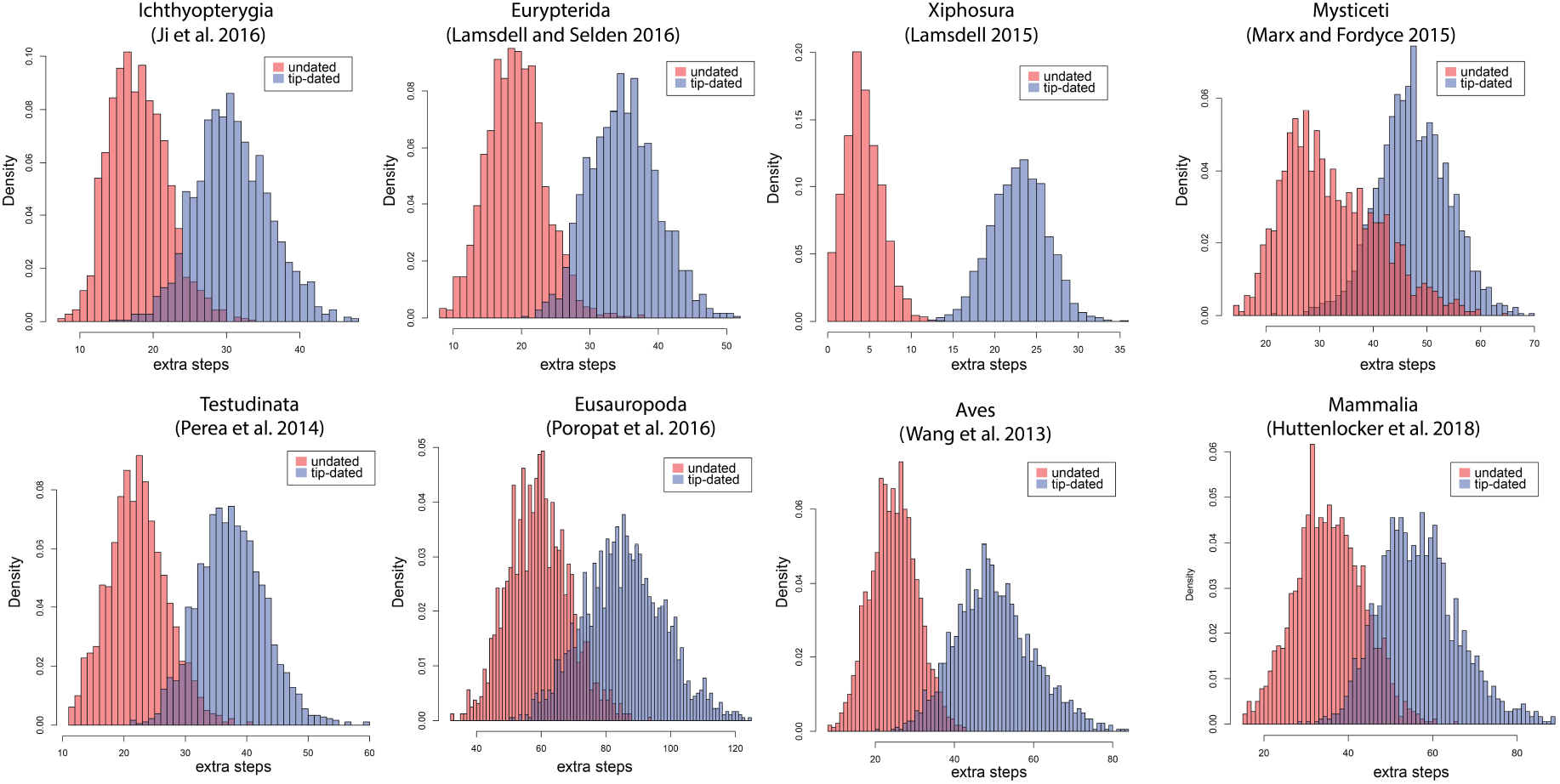
Bayesian tip-dating produces trees that are less parsimonious than undated Bayesian analysis. Histograms of the number of additional steps required by trees produced by tip-dating and undated Bayesian analysis, compared to the tree length of the most parsimonious trees. For the smallest dataset (Xiphosura), some of the trees produced by undated analysis are identical to parsimony trees.

### Stratigraphic Fit

P-values for all stratigraphic fit calculations were highly significant, showing that all methods produce trees with better fit to stratigraphy than expected by chance. As expected, tip-dating approaches produce trees with a better stratigraphic fit than undated Bayesian or parsimony (Fig. 3). Trees produced when the analysis samples solely from the prior have a particularly high stratigraphic fit, again as expected. Plots showing the prior probability of trees against stratigraphic fit for each dataset show positive and highly significant correlations across all methods (Fig. 3). Results for the Stratigraphic Completeness Index (Fig. S2) are similar to those for the Gap Excess Ratio (Fig. 3), with the exception of the sauropods dataset, which shows no positive correlation for the SCI.

**Figure 3.**
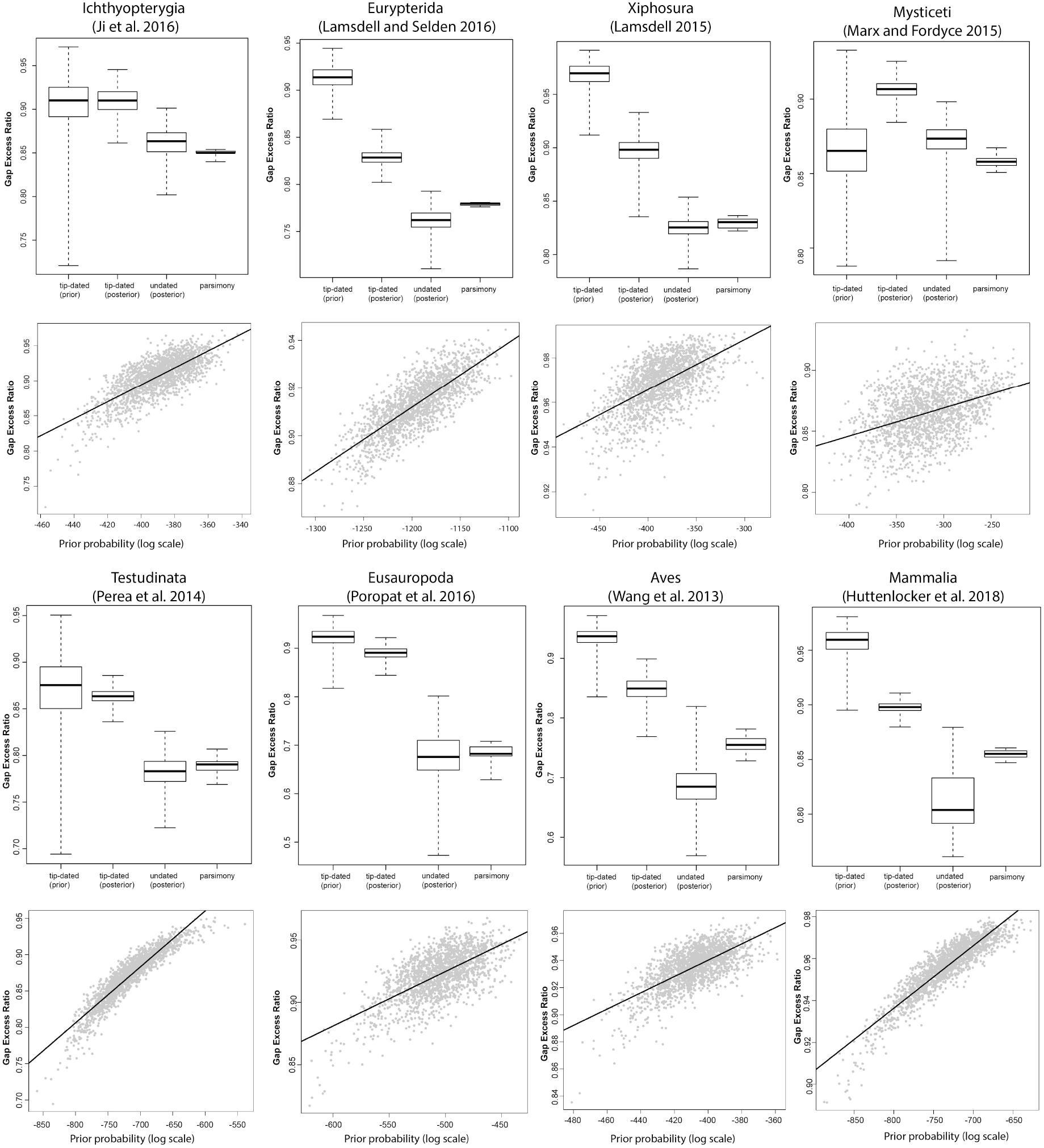
Bayesian tip-dating methods recover trees with better fit to stratigraphy than other methods. Upper panels: Gap Excess Ratio for every tree in each sample is shown as a box plot (whiskers span full range). Lower panels: A positive correlation exists between fit to stratigraphy and prior probability for every dataset (each data point represents a tree from the prior sample for tip-dating).

Stratigraphic fit is lower for the posterior sample of trees from the tip-dating analysis compared to the prior, but still higher than trees for the other two methods. There is little difference between the stratigraphic fit measures for the parsimony and undated Bayesian methods relative to the differences with the tip-dated results. However, we note that Sansom et al. (2018) reported a small but statistically significant difference favouring parsimony based on many datasets (although results vary within particular datasets). Our results suggest that tip-dating methods assign a higher prior probability to trees with a better stratigraphic fit, as expected. This leads to a better stratigraphic fit for the tree topologies in the posterior sample, compared to trees produced by other methods. This is likely to be the cause of the topology differences between tip-dating and the other methods shown in the previous section.

### Effect of Incomplete Taxa

A feature of Bayesian analyses is that the prior is most important when data are weak, while strong data overwhelm the prior (Edwards et al. 1963). Since tip-dating places a higher prior probability on trees with better fit to stratigraphy, it is reasonable to expect that this becomes most important in parts of the phylogeny that are only weakly or incompletely resolved by morphological data. Iterative deletion of incomplete taxa from the sauropods dataset supports the hypothesis that much of the difference in tree topology and stratigraphic fit between tip-dated and undated Bayesian methods are in parts of the phylogeny which cannot be confidently resolved by the morphological data. With each successive deletion of incomplete taxa, the stratigraphic fit of trees from the undated Bayesian analyses increases, whereas the stratigraphic fit of the tree from tip-dating is essentially unchanged (Fig. 4a). Topological differences between the methods generally decline with each deletion (Fig. 4b). Random deletion of taxa does not lead to changes in stratigraphic fit for either method (Fig.4c), and topological differences do not change (Fig. 4d). This shows that the observed patterns are due to the deletion of incomplete taxa, not merely a result of deletion of taxa in general. These results suggest that tip-dated analyses constrain the phylogenetic position of incomplete taxa based on their stratigraphic age, leading to topological differences when compared to other methods. Parsimony analyses were also performed on the same datasets, but the results did not produce any pattern, likely due to the very low number of trees found for most datasets.

**Figure 4.**
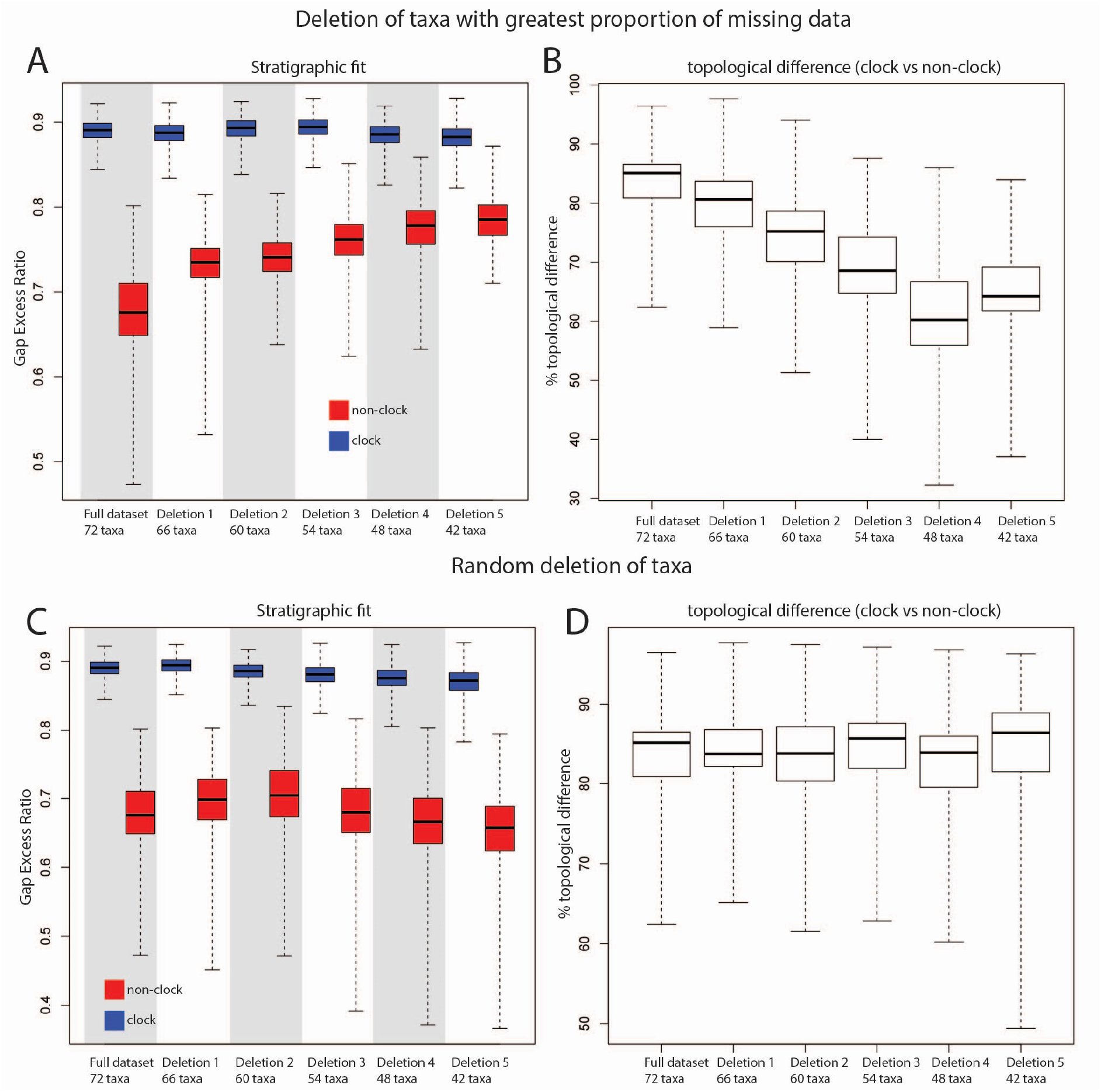
Differences between tip-dating and undated Bayesian methods in terms of topology and stratigraphic fit are concentrated in uncertain parts of the phylogeny. This study utilises the sauropod dataset of Poropat et al. (2016). Successive deletion of the most incomplete taxa (top) leads to an increase in stratigraphic fit for undated analysis, but not tip-dating. Topological differences between these methods also successively decrease. Random deletion of taxa (bottom) shows that these patterns are not purely an artefact of fewer taxa.

### Mesozoic Mammals and the Allotheria Hypothesis

Tip-dating analysis of the complete Mesozoic mammal dataset (Fig. 5) resulted in taxa that have been previously been suggested to belong to Allotheria falling into three separate clades. The late Triassic haramiyidans *Haramiyavia* and *Thomasia* are placed outside Mammaliaformes, in a strongly-supported clade with tritylodontids (BPP = 0.97). The middle Jurassic euharamiyidans, early Cretaceous hahnodontids, and the late Cretaceous Madagascan gondwanatherian *Vintana*, by contrast, collectively form a strongly-supported clade (BPP =1.00) within Mammaliaformes, although our phylogeny is insufficiently well-resolved to indicate whether or not this is within crown-clade Mammalia. As in Huttenlocker et al. (2018), *Vintana* is nested within a clade that also includes euharamiyidans and the hahnodontids *Hahnodon* and *Cifelliodon*, and is sister to *Cifelliodon*. Finally, the multituberculates form a third strongly-supported clade (BPP = 1.00), within Mammalia, that falls closer to Theria than to Monotremata. Polyphyly of Allotheria has been recovered in several previous analyses (e.g. Gurovich and Beck 2009, Luo et al. 2015, Huttenlocker et al. 2018), whereas others have recovered a paraphyletic Haramiyida within a monophyletic Allotheria (Bi et al. 2014, Krause et al. 2014), but this is the first time (to our knowledge) that a formal phylogenetic analysis has recovered polyphyly of Haramiyida in addition to allotherian polyphyly.

**Figure 5.**
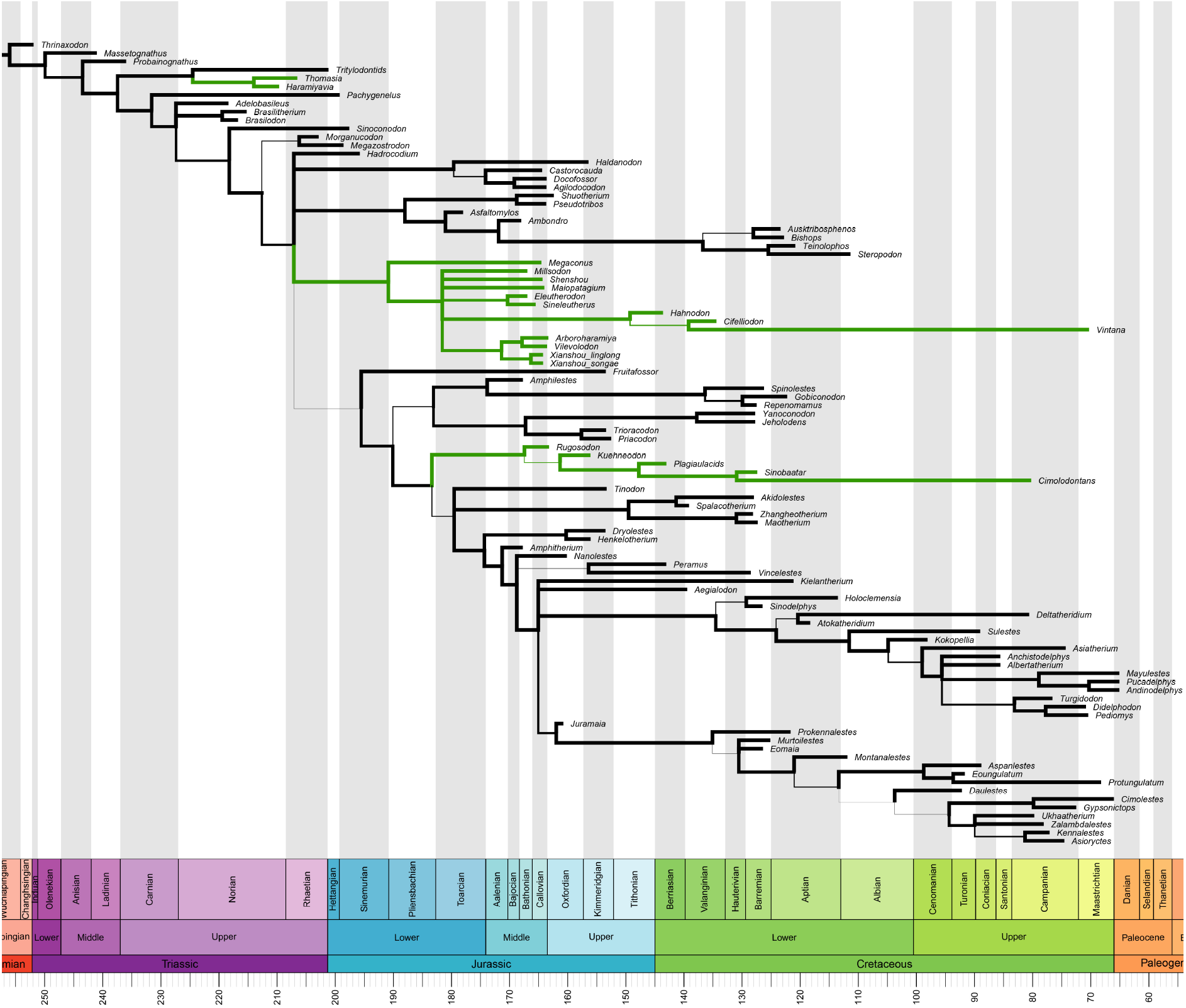
50% majority rule consensus tree for Mesozoic mammals from a tip-dated analysis of the dataset in Huttenlocker et al. (2018). So-called allotherians (*Haramiyavia, Thomasia*, euharamiyidans, multituberculates, hahnodontids and the gondwanatherian *Vintana*) are in green. Branch widths proportional to posterior probability (between 0.5 and 1.0).

**Figure 6.**
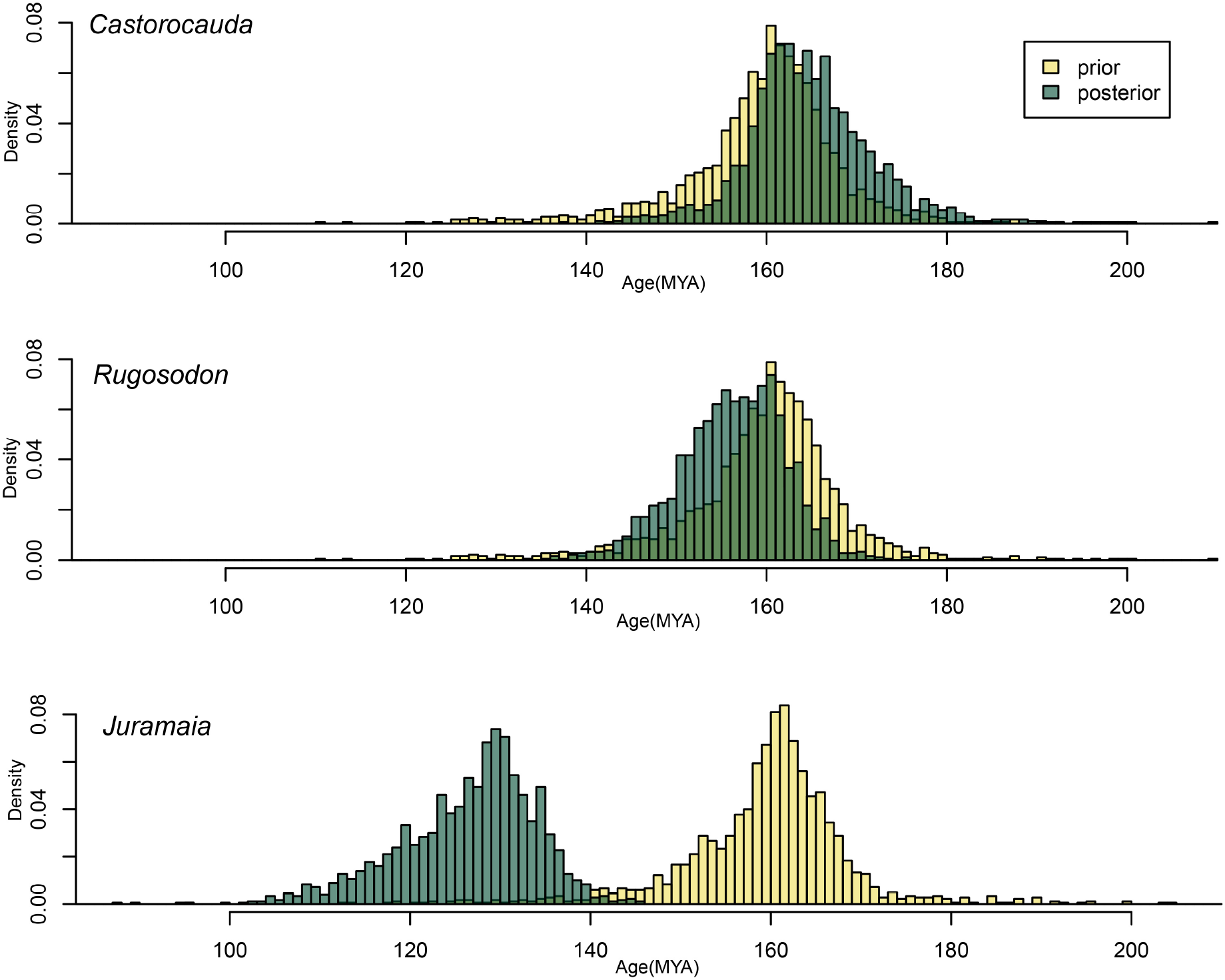
The morphological data has a strong signal towards a Lower Creataceous rather than a jurassic age for *Juramaia*. Age estimates for 3 putative members of the Yanliao biota, analysed with a Laplace distribution prior centred on 161 MA, therefore representing a conservative test for the age of *Juramaia*. Data for *Castorocauda* and *Rugosodon* place these taxa firmly in the correct age range for the Yanliao biota, in contrast to the data for *Juramaia* which strongly contradicts the prior.

Support for monophyly of Allotheria and of Haramyida appears to be driven by dental characters: when we repeated our tip-dating analysis using dental characters only, we recovered a strongly-supported clade comprising tritylodontids, haramyids, euharamyidans, hahnodontids, *Vintana* and multituberculates, which falls outside Mammaliaformes (Fig. S3). However, when we analysed cranial and dental characters together, we recovered approximately the same relationships as in our complete analysis (which also includes postcranial characters), with “allotherians” split between three separate clades (Fig. S4). Tip-dating analysis of postcranial characters only also recovers separate euharamyidan and multituberculate clades (Fig. S5), but *Haramiyavia, Thomasia*, hahnodontids and *Vintana* could not be included in this analysis as postcranial remains have not been described for them (undescribed postcranial material is known for *Haramiyavia*) (Jenkins et al. 1997, Luo et al. 2015)

Tip-dating using the dataset of Krause et al. (2014) led to a more complex result, as the sample of post-burn trees include topologies in which Allotheria is polyphyletic and in which it is monophyletic This analysis required extremely long run times over an unusually large number of independent runs to reach convergence due to the “twin peak” behaviour displayed by the prior and the likelihood traces. These peaks correspond to the two different tree topologies regarding Allotheria. One peak, where *Haramiyavia* and *Thomasia* formed a clade with other allotherians (essentially the parsimony result) had a low prior probability but high likelihood (Fig. S6A). The other peak, which had *Haramiyavia* and *Thomasia* closer to the root of the tree, and separated from other allotherians, had a higher prior probability and lower likelihood (Fig. S6B). Overall, polyphyly of Allotheria (with *Haramiyavia* and *Thomasia* forming a separate clade closer to the root) was the preferred hypothesis, found in 63% of the posterior sample (Figs. S7, S8) compared with 37% showing monohyly of Allotheria (Fig. S9). The position of the enigmatic gondwanatherians was highly labile, and they were excluded from the trees for these calculations. *Arboroharamiyavia*, the only euharamiyidan included in the Krause et al. (2014) dataset was always recovered in a clade with multituberculates. The Allotheria polyphyly hypothesis had only a slight advantage in terms of posterior probability (Fig. S6C) relative to the Allotheria monophyly hypothesis, as the likelihood and prior probabilities almost cancel out. A constrained parsimony search revealed that polyphyly of Allotheria requires four additional steps (a 2.23% increase in treelength) compared to the unconstrained analysis (which recovers allotherian monophyly). However, constrained and unconstrained trees were not significantly different (p=0.68) under the Templeton test (Templeton 1983).

### The Problem of Juramaia

Rerunning the analysis with a Laplace prior distribution on the age of putative Middle Jurassic eutherian *Juramaia*, centred on 161 MYA but without a hard upper or lower bound had no effect on the recovered relationships of haramiyidans and multituberculates: haramiyidan diphyly (and allotherian triphyly) was still recovered (Fig. S10). Strikingly, however, this analysis revealed a strong signal in the data supporting a younger age for *Juramaia*. The mean estimated age for *Juramaia* was 126.4 Ma (interestingly, almost exactly the same as the age of the Yixian Formation, from where several fossil eutherians are known, such as *Eomaia* and *Ambolestes)(Bi* et al. 2018), with a 95% highest posterior density (HPD) interval of 111.4 – 140.2 Ma. Thus, the 95% HPD is entirely within the Early Cretaceous. This contrasts with the results from *Castorocauda* and *Rugosodon*, uncontroversial members of the Middle Jurassic Yanliao Biota, when these were assigned the same Laplace prior age distributions as *Juramaia*. The mean age estimate for *Rugosodon*, which is from the slightly younger (164-159 MYA) Linglongta subdivision of the Yanliao Biota, was 156.2 Ma (HPD 144.0 – 166.7). That for *Castorocauda*, which is from the slightly older (168-164 MYA) Daohugou subdivision, was 164.6 Ma (HPD 150.4 – 179.5). The estimated ages of the Linglongta and Daohugou Biotas are therefore included in the 95% HPDs for *Rugosodon* and *Castorocauda* respectively.

### Prior Probabilities on Tree Topology for Haramiyidans

The results show that the prior for the placement of the Middle Jurassic euharamiyidans (Fig. 7a) is very different from that of the late Triassic *Haramiyavia* and *Thomasia* (Fig. 7b). The prior for *Haramiyavia* and *Thomasia* is concentrated around the very oldest part of the tree: the time at which *Haramiyavia* and *Thomasia* branch from the rest of the tree has a highest prior density interval of 205 – 242.4 Ma. The prior for euharamiyidans is concentrated on younger branches (HPD 173.2 – 217 Ma). Notably, both groups are also more likely to attach to longer branches (Fig. 7). When the prior is corrected for the effect of branch length, by dividing prior probability by branch length (Fig. S11), the temporal signal is more obvious. Quantification of the prior probability that each clade occurs above a particular node also shows that the prior for *Haramiyavia* and *Thomasia* is strongly concentrated at the very base of the tree, in contrast to the prior for euharamiyidans (Fig. 7c).

**Figure 7.**
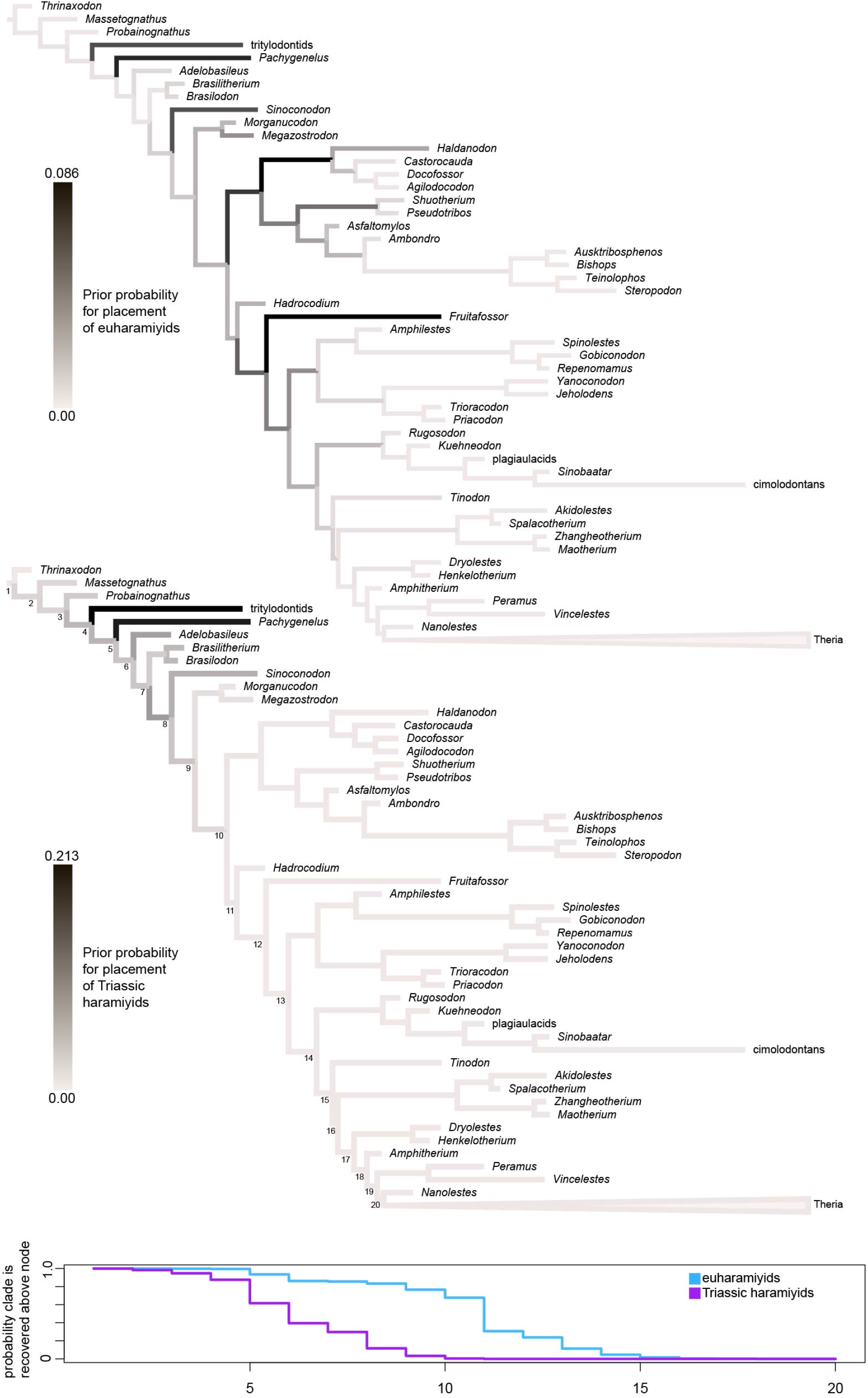
Topology prior for haramiyidans. A-B) The tree is a fixed topology based on the maximum clade credibility tree from the main analysis, on which the prior probabilities for the position of two groups of haramiyidans are mapped. Branch colours represent the prior probability that the respective clade (A, Euharamiyida; B, *Haramiyavia*+*Thomasia*) was found on that branch in an analysis run without data. The prior probabilities for a clade comprising *Haramiyavia* and *Thomasia* (both of which are late Triassic) are concentrated at the base of the tree, unlike the prior for euharamiyidans (which are all middle Jurassic), which is more diffuse and centred in younger parts of the tree. There is also a notable prior preference for long branches. C) Prior probabilities that each group of haramiyidans is found above each node in a sequence from the base of the tree to the crown therian node. The distribution for *Haramiyavia+Thomasia* drops off much more quickly, representing the prior expectation that they are placed in a lower position on the tree than euharamyidans.

This analysis is only an approximation of the true prior, as in reality the topology of the whole tree is also subject to change. However, it shows that the analysis has a prior expectation that *Haramiyavia* and *Thomasia* are separate from euharamiyidans, due to the differing ages of the two sets of taxa. Parsimony analysis with a negative constraint on haramiyidan monophyly (i.e. preventing *Haramiyavia* and *Thomasia* forming a clade with euharamiyidans) produce trees that are only two steps longer (representing just a 0.1% increase in tree length) than the unconstrained trees. Constrained and unconstrained trees were not significantly different (p=0.87) under the Templeton test (Templeton 1983). This is likely to be a factor in the recovery of haramiyidan diphyly in the posterior, as the morphological character data only weakly supports haramiyidan monophyly, whereas the (temporally-influenced) prior strongly supports diphyly.

## Discussion

### Differences in Tree Topology between Different Methods

The question of whether parsimony or Bayesian methods should be used for phylogenetic analysis of morphology-only datasets has been the subject of a vigorous debate (Wright and Hillis 2014, O’Reilly et al. 2016, Goloboff et al. 2017, Goloboff et al. 2018, O’Reilly et al. 2018, Sansom et al. 2018). These studies have focused on undated Bayesian approaches, with tip-dating receiving relatively little attention. However, our results show that tree topologies produced by tip-dating are outliers when compared to trees produced by parsimony and undated Bayesian methods, a result also found for one dataset in Turner et al. (2017).

Much of the debate between parsimony and undated Bayesian methods has centred on consensus trees and their degree of resolution (O’Reilly et al. 2016, Brown et al. 2017, Goloboff et al. 2017). Our results comparing topological differences within methods show that parsimony is (with one exception) more resolved (Fig. 1, Fig. S1), as previously reported (O’Reilly et al. 2016). Bayesian tip-dating and undated methods are in general approximately equally resolved (Fig.1, Fig. S1). The topological differences between tip-dating and other methods are therefore likely driven largely by the incorporation of stratigraphic ages and the birth-death-sampling model, with the use of the Mkv model a relatively minor cause of the topological differences between methods.

### When will Stratigraphic Data Overrule Morphological Data?

The difference in tree topology appears to be driven by the effective prior probabilities placed on tree topologies in the tip-dating analysis. Tree topologies with a better stratigraphic fit (i.e. requiring less unsampled branch length, or shorter ghost lineages) are given a higher prior probability (Fig. 3). The stratigraphic fit of the posterior distribution of tree topologies is intermediate between the prior and the values from undated Bayesian and parsimony approaches (Fig. 3), reflecting the interplay of the prior and the evidence from the morphological character data.

The interplay of the prior and the likelihood is demonstrated well in the analyses of the Mesozoic mammal datasets of Huttenlocker et al. (2018) and Krause et al. (2014). Tip-dating of the Huttenlocker et al. (2018) dataset results in strong support for polyphyly of Allotheria, including diphyly of the haramiyidans, and haramiyidan diphyly requires only two additional steps under parsimony. In contrast, the analysis of the Krause et al. (2014) dataset flips between allotherian polyphyly and monophyly, and allotherian polyphyly requires 4 additional parsimony steps over monophyly. In the case of the Krause et al. (2014) dataset, the stronger morphological signal for allotherian monophyly is therefore not fully overruled by the stratigraphic evidence incorporated into the tree prior.

In Bayesian analysis, the prior becomes more important when the data are weak, so it might be predicted that the prior probabilities favouring trees with good fit to stratigraphy are most influential in weakly resolved parts of the tree. This was tested by successive deletion of the most incomplete taxa from the sauropods dataset (Fig. 4). As hypothesised, this led to successive increases in the stratigraphic fit of topologies estimated by undated Bayesian analyses, and a decrease in the topological differences between undated and tip-dating Bayesian approaches (no changes occurred when taxa were deleted randomly with respect to completeness). This suggests that much of the topological difference between tip-dating and other phylogenetic methods is driven by the placement of incomplete taxa. In undated analyses (under either parsimony or model-based optimality criteria), these incomplete taxa may fit into several positions on the tree even if these are incongruent with their stratigraphic age, but such stratigraphically incongruous positions require stronger morphological evidence in the tip-dating approach. These results suggest that, as morphological data for incomplete taxa improve, topologies recovered from undated Bayesian or parsimony approaches might become more similar to those produced by tip-dating. This is echoed in the results of Benton and Storrs (1994), which showed that phylogenetic trees produced in 1993 generally had a higher stratigraphic fit than comparable trees from 1967.

Amongst the datasets investigated, stratigraphic fit measures are lowest in the eurypterid dataset, aligning with findings that, in general, stratigraphic fit for arthropod phylogenies are lower than for other groups (O’Connor and Wills 2016). The oldest eurypterid, the Middle Ordovician *Pentecopterus*, is found in a deeply nested position, together with Late Ordovician members of the Megalograptidae, with both parsimony analysis (Lamsdell and Selden 2017) and tip-dating (Fig. S12). This shows that even highly stratigraphically incongruent topologies can still be recovered if there is sufficient morphological evidence: the branch leading to Megalograptidae (including *Pentecopterus*) has five unambiguous character changes. In addition, three of the branches between the root and the megalograptid clade are supported by at least five unambiguous synapomorphies. Notably, the tip-dating analysis for eurypterids also estimated an ancient divergence date for eurypterids, more than 30 million years before the appearance of *Pentecopterus*. A younger divergence date would require extremely rapid divergences at the base of tree, violating the assumptions of relatively constant diversification rates in the tree prior.

These results support the view that inclusion of stratigraphic age data in tip-dated Bayesian phylogenetic analysis directly affects tree topology. Highly incomplete taxa are more constrained to positions congruent with stratigraphy in tip-dated analysis, even if the (limited) character data are also consistent with other, less stratigraphically congruent positions. Conversely, taxa for which abundant data are available can be placed in stratigraphically incongruent positions if there is sufficient character support. This is intuitive, and in some respect already resembles the approach already informally taken by most palaeontologists: extraordinary claims require extraordinary evidence.

### Is the Inclusion of Stratigraphic Data in Phylogenetic Analysis Justified?

> “It is merely eccentric to claim that time is not a desirable parameter in working out phylogenies, even though what is desirable is not invariably available or clear in its significance” (Simpson 1975).

Fisher (2008) raised arguments against criticisms of stratocladistics, including those regarding the imperfect nature of the fossil record, and these will not be repeated here. The fossil record need only be adequate, not perfect, to justify its utilisation (Paul 1985). The high level of congruence between phylogenetic and stratigraphic data (Gauthier et al. 1988, Norell and Novacek 1992, Benton and Hitchin 1997) shows that stratigraphic data is generally informative regarding clade age and branching order. Including both stratigraphic and morphological data when estimating phylogeny allows these two generally congruent data types to reciprocally illuminate in the minority of occasions that they disagree.

Stratigraphic data will always be incomplete (in the sense that not every species will be present in the fossil record), but the same can be said of morphological data. No fossil preserves 100% of morphological data (even a fossil being scored for 100% of characters within a particular morphological data matrix is rare). In addition, convergent evolution means that morphological data can sometimes be misleading, a fact becoming increasingly apparent with the availability of increasing amounts of molecular data (Lee and Palci 2015). Stratigraphic data provides an opportunity to identify probable convergent evolution in clades widely separated by time, a point that has already been demonstrated in tip-dated phylogenetic analyses of palaeognath birds (Worthy et al. 2017) and gharials (Lee and Yates 2018).

Stratigraphic and morphological data represent a way of assessing the quality of each other. Stratigraphic incongruence cannot be recognised independently from morphological data, but then how do we know that the morphological data are correct? In the absence of molecular data, stratigraphic data are the only independent assessment of the quality of morphological data. Simultaneous analysis of both forms of data is therefore a good approach, and should avoid the recovery of highly stratigraphically incongruent trees based on scant evidence. However, morphological data have generally been afforded precedence. For example, Turner et al. (2017) suggest, based on analysis of a crocodylomorphs dataset, that tip-dating should not be used in cases where sampling is uneven, leading to long un-sampled branches which are then disfavoured by tip-dating. However, their recognition of uneven sampling was not independent from phylogeny, and was based on a time-scaled parsimony tree of the same dataset. On the contrary, an expectation that long un-sampled branches should be rare is a reasonable prior, one that requires solid data to overturn.

### Mesozoic Mammals and Convergent Dental Evolution

Marsh (1880) erected the order Allotheria for two multituberculate taxa, *Ctenacodon* and *Plagiaulax*, and placed Allotheria within Marsupialia. Since then, a number of different fossil taxa that share a similar, superficially rodent-like dentition (most notably, postcanine teeth with cusps in longitudinal rows) have been referred to Allotheria at one time or another, namely: tritylodontids (now recognised to be non-mammaliform cynodonts), multituberculates, haramiyidans and gondwanatherians (Kielan-Jaworowska et al. 2004). However, there has long been controversy as to whether Allotheria is monophyletic or not, with most of the recent debate centred on whether haramiyidans form a clade with multituberculates (Gurovich and Beck 2009, Bi et al. 2014, Krause et al. 2014, Luo et al. 2015, Chang et al. 2017, Huttenlocker et al. 2018, Meng et al. 2018). Also controversial has been the position of these so-called allotherian taxa (regardless of whether Allotheria itself is monophyletic) within broader mammaliaform phylogeny. Although it has long been recognised that so-called allotherians are not closely related to marsupials (as originally proposed by Marsh, 1880) or other therian mammals, exactly where they fit relative to crown-clade Mammalia has remained uncertain. Resolution of this issue is key for dating the origin of Mammalia: if the late Triassic haramiyidans *Haramiyavia* and *Thomasia* fall within Mammalia, then the divergence between monotremes and therians is at least this old (Zheng et al. 2013, Bi et al. 2014). It also has implications for the evolution of key mammal features. Perhaps most significantly, if haramiyidans do not form a clade with multituberculates, then it implies that these two lineages independently acquired the definitive mammalian middle ear (Meng et al. 2018). Finally, there are also implications for the origin of the enigmatic gondwanatherians, specifically whether or not this group is related to multituberculates (Gurovich and Beck 2009, Krause et al. 2014).

The results of our tip-dating analyses indicate that the dental similarities proposed to unite Allotheria are homoplastic, and that they evolved at least three times independently: once in the common ancestor of *Haramiyavia+Thomasia* and tritylodontids, once in the common ancestor of euharamiyidans, hahnodontids and gondwanatherians, and once in multituberculates. The finding that the dental resemblances between allotherians is the result of convergence is perhaps unsurprising, as broadly similar combinations of dental features has also evolved in placental mammals (e.g. rodents) and in metatherians (e.g. polydolopimorphians). More generally, one recent study found that dental characters in mammals are more prone to homoplasy than characters from the rest of the skeleton (Sansom et al. 2017). In our tip-dating analyses, evidence from the cranium and (for those taxa known from postcranial remains) postcranium, together with temporal information, is sufficient to overwhelm the dental signal that favours haramiyidan and allotherian monophyly.

Perhaps the most novel aspect of our results is diphyly of Haramiyida: the late Triassic *Haramiyavia* and *Thomasia* form a clade with tritylodontids outside Mammaliformes, distinct from a separate clade comprising the euharamiyidans from the middle Jurassic of China, the early Cretaceous hahnodontids *Hahnodon* and *Cifelliodon*, and the Madagascan Late Cretaceous gondwanatherian *Vintana. Haramiyavia* and *Thomasia* have been placed with tritylodontids in some previous analyses (e.g. Gurovich and Beck 2009), but it should be noted that these two groups show marked dental differences (Kemp 1982, Jenkins et al. 1997, Kielan-Jaworowska et al. 2004, Luo et al. 2004, Kemp 2005, Luo et al. 2015, Velazco et al. 2017): the upper postcanines of tritylodontids comprise three major rows of cusps arranged labiolingually, whereas those of *Haramiyavia* and *Thomasia* have only two; the dental power stroke is strongly palinal in tritylodontids, whereas it is a predominantly orthal in *Haramiyavia* and *Thomasia*; tritylodontids show a specialised type of postcanine dental replacement, in which worn teeth shed from the anterior end of the toothrow and new teeth are added at the posterior end, whereas this is absent in *Haramiyavia* at least (the pattern of dental replacement in *Thomasia* is unknown); *Haramiyavia* also retains the upper and lower canines, whilst these teeth have been lost by tritylodontids. Based on these striking morphological differences, this relationship should be viewed cautiously pending description of additional cranial and postcranial material of *Haramiyavia, Thomasia* or related taxa that might corroborate it. This relationship may also partly be driven by the prior in our analysis (Fig. 7) due to the relatively long branch leading to tritylodontids, which also means that the morphological differences between tritylodontids and *Haramiyavia* and *Thomasia* will be penalised less.

Diphyly of Haramiyida is perhaps surprising, and has not been found in previous analyses. However (among other differences) the best known late Triassic haramiyidan, *Haramiyavia*, retains a prominent postdentary trough (Luo et al. 2015), a plesiomorphic feature indicating that it lacked a definitive mammalian middle ear, whereas in most euharamiyidans (with the notable exception of *Vilevolodon*; (Luo et al. 2017)) this trough is either very small or absent (Zheng et al. 2013, Bi et al. 2014, Chang et al. 2017, Meng et al. 2018). Ongoing research has revealed previously unsuspected levels of homoplasy in the evolution of the DMME throughout mammaliaform phylogeny, and current evidence indicates that this supposedly “key” mammalian feature arose multiple times (Luo 2007, Luo 2011, Bi et al. 2014, Meng 2014, Ramírez-Chaves et al. 2016, Chang et al. 2017, Urban et al. 2017, Meng et al. 2018). Nevertheless, the incorporation of temporal information in our tip-dating analysis is sufficient to split the middle Jurassic euharamiydans from the late Triassic *Haramiyavia* and *Thomasia*, indicating the comparatively weak morphological character support for monophyly of Haramiyida.

### Age of Juramaia

The holotype and only known specimen of *Juramaia sinensis* (BMNH3413) is reported as having been collected from the Daxigou (or Daxishan) locality of Jianchang County, Liaoning Province, China, within the Lanqi/Tiaojishan Formation (Luo et al. 2011, Xu et al. 2016). Radiometric dates at this locality suggest that the fossil-bearing beds at this locality (which preserve the Linglongta Biota, the younger of the two subdivisions within the Yanliao Biota) are 164-159 million years old (Xu et al. 2016). However, several papers have raised questions about the age of *Juramaia*, based largely on its comparatively derived dental morphology, which is very similar to eutherians that are about 35 million years younger such as *Eomaia* and *Ambolestes* from the Yixian Formation (Meng 2014, Sullivan et al. 2014, Bi et al. 2018). Indeed, based on this resemblance, Sullivan et al. (2014) specifically raised the possibility that *Juramaia* might be from a Cretaceous site elsewhere in Jianchang County. Confident resolution of this issue will probably require more detailed studies of the holotype, and/or discovery of additional *Juramaia* specimens at Daxigou/Daxishan. However, the results of our tip-dating analyses when the age of *Juramaia* is allowed to vary are striking: a mean estimate of 126.4 MYA, almost identical to the age of *Eomaia* and *Ambolestes* (Chang et al. 2017, Bi et al. 2018), with the age of Daxigou/Daxishan exceeding the upper bound of the 95% HPD by ~20 Ma. Also notable is the fact that the estimated ages of two other mammaliforms from the Yanliao Biota, *Castorocauda* and *Rugosodon* remain centred on the middle Jurassic when their ages were allowed to vary. For some datasets at least, Bayesian tip-dating appears to perform relatively well at estimating the ages of tips when treated as unknown (Drummond and Stadler 2016), although 95% HPDs can be wide (Dembo et al. 2016). Thus, our results provide at least circumstantial evidence that the age of *Juramaia* may be younger than currently interpreted. If so, then the palaeontological minimum for the timing of the split between Eutheria and Metatheria is shifted forward to the early Cretaceous, either 145 MYA (if the recently described but poorly preserved *Durlstotherium* and *Durlstodon* from southern England are eutherians; (Sweetman et al. 2017) or 126 MYA (the age of the well-preserved Yixian eutherians (Chang et al. 2017, Bi et al. 2018).

## Acknowledgements

We thank Mike Lee, John Long, Matt Friedman and Martin Brazeau for comments on an earlier version of the manuscript, and Graeme Lloyd for discussions. BK thanks Martin Rücklin for generously supporting the completion of this paper. Tnt is made freely available through the Willi Hennig society.

