## Supplementary material for "Bayesian Tip-dated Phylogenetics: Topological Effects, Stratigraphic Fit and the Early Evolution of Mammals": Fig. S

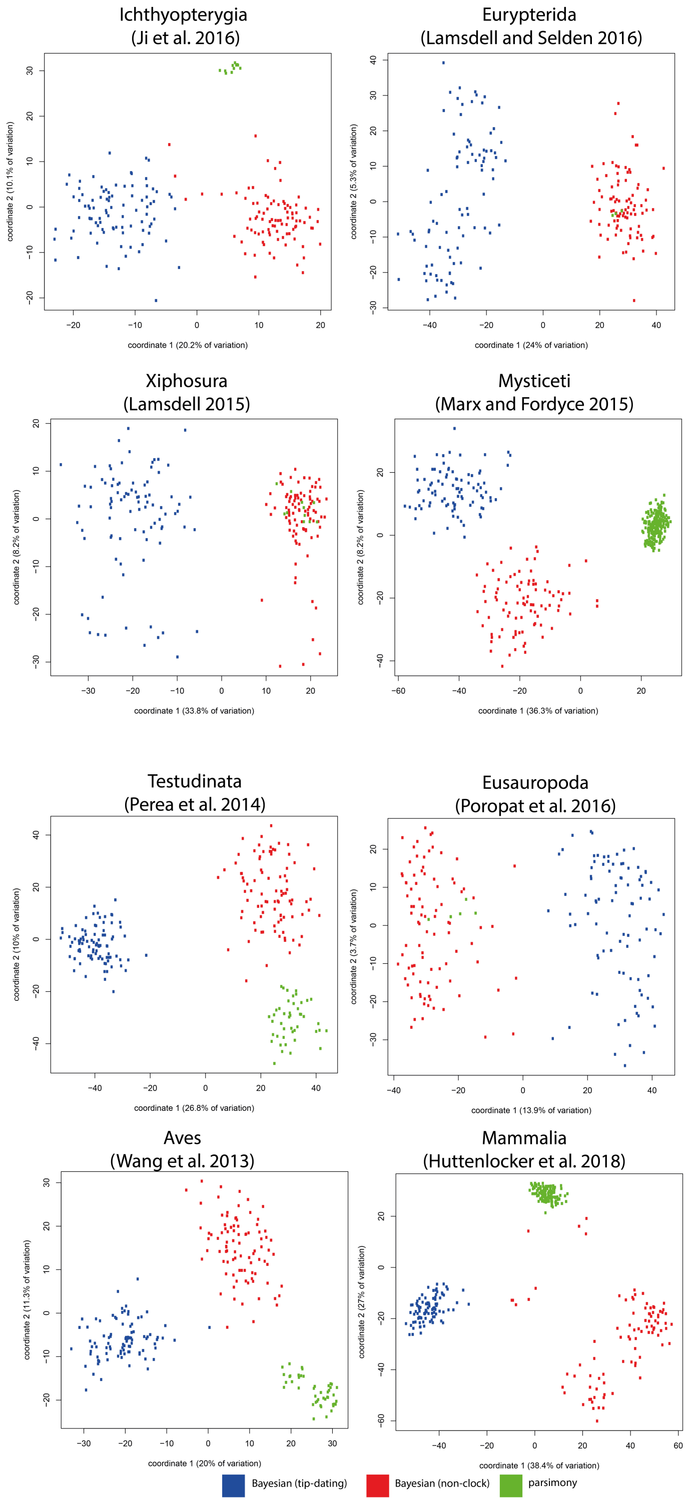


**Figure S1. Topological differences between different phylogenetic methods (tip-dated Bayesian, undated Bayesian and parsimony) as visualised through principal coordinates plots of Robinson-Foulds distances.** Overall, parsimony and undated Bayesian methods are similar, and plot in a single cluster in 3 of the 8 datasets.

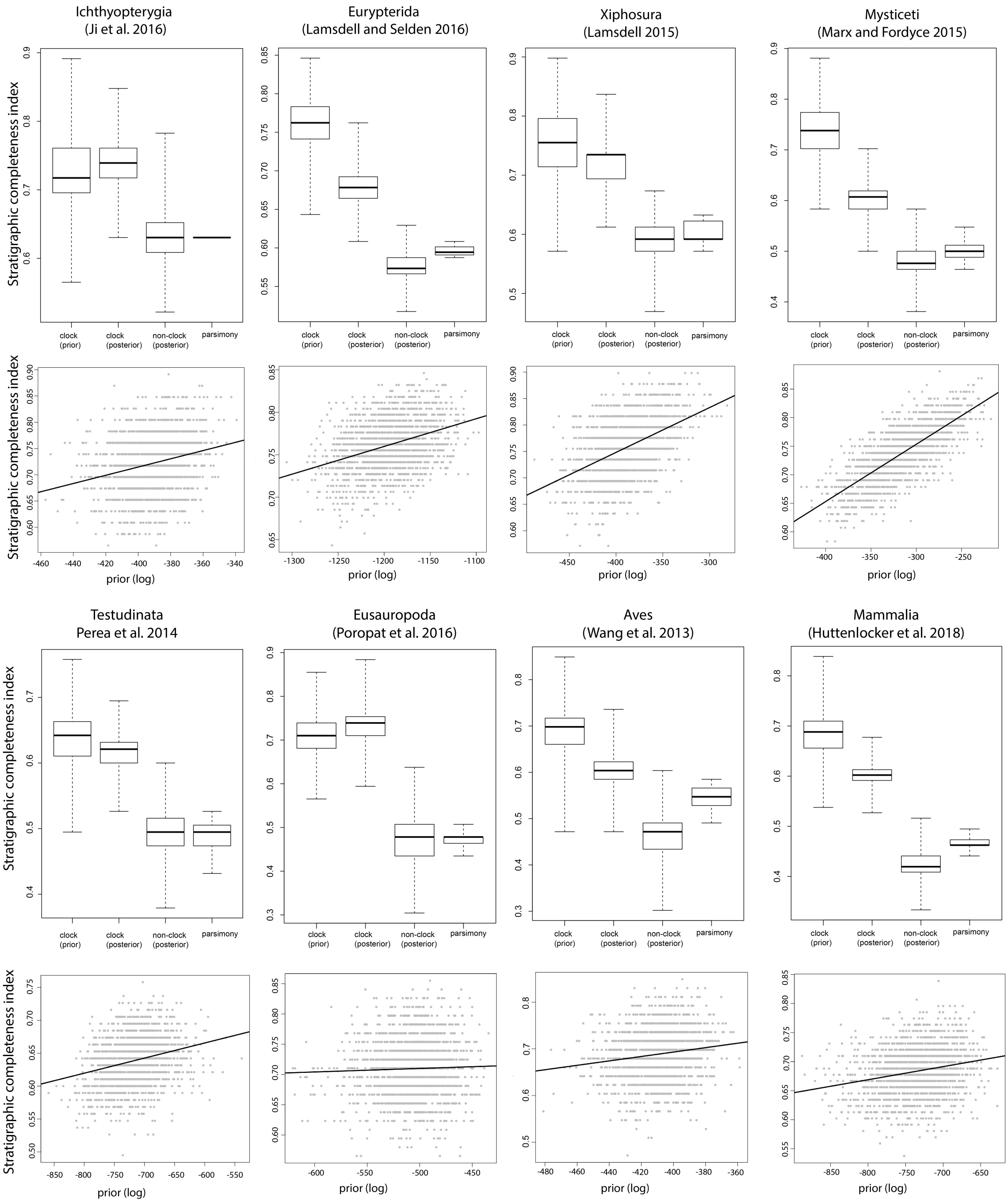


**Figure S2. Bayesian tip-dating methods recover trees with better fit to stratigraphy than other methods.** Upper panels: Stratigraphic consistency index for every tree in each sample is shown as a box plot (whiskers span full range). Lower panels: A positive correlation exists between fit to stratigraphy and prior probability for every dataset except sauropods (each data point represents a tree from the prior sample for tip-dating).

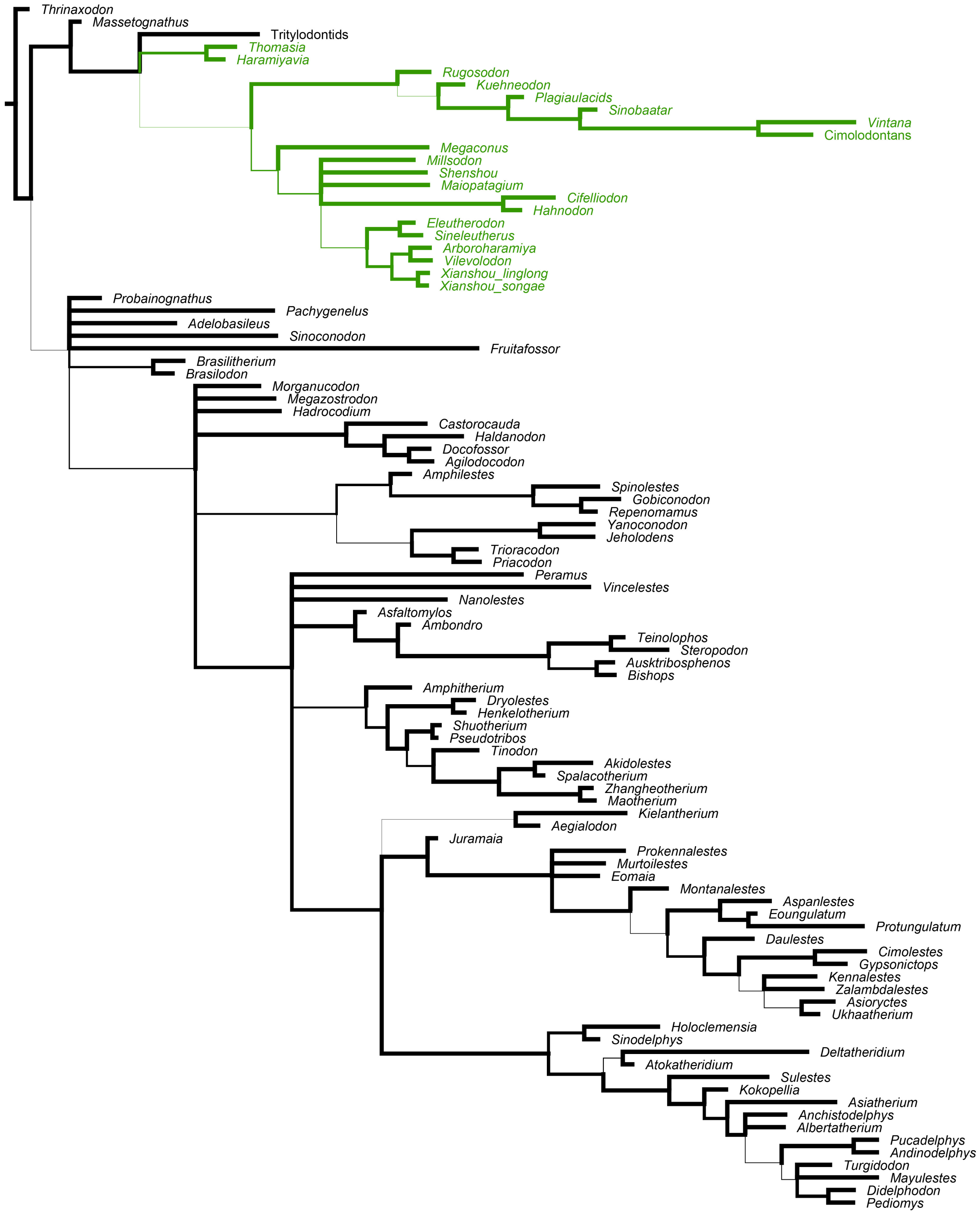


**Figure S3. Majority rule consensus tree of tip-dated analysis of dental characters from the dataset of Huttenlocker et al (2018).** Here the allotherians (highlighted in green) are recovered as a monophyletic clade outside crown Mammalia. Branch widths proportional to support (between 0.5 and 1.0).

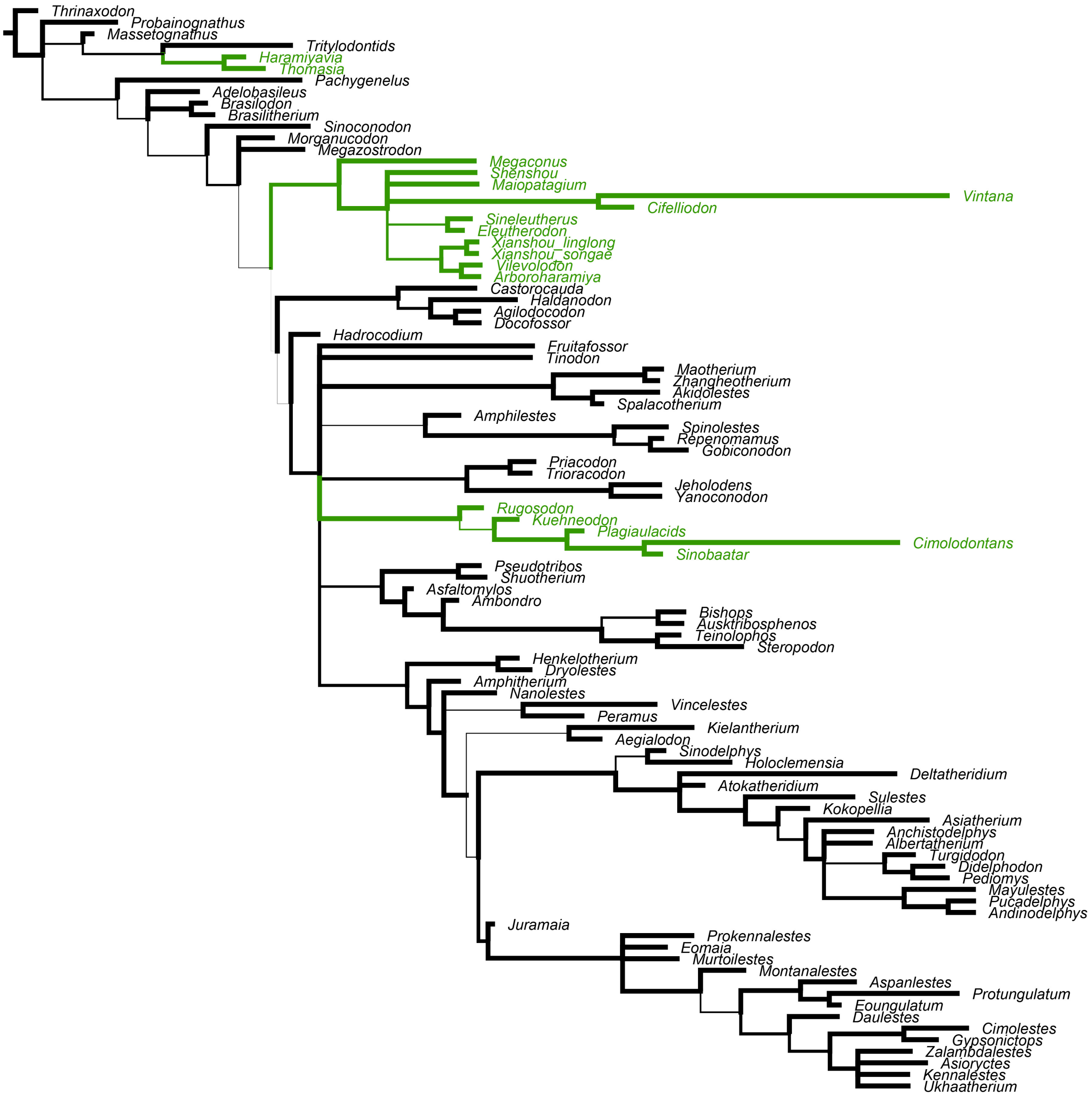


**Figure S4. Majority rule consensus tree of tip-dated analysis of cranial and dental characters from the dataset of Huttenlocker et al (2018).** This phylogeny is broadly congruent with the results of the analysis of the complete dataset. Allotherians (green) are recovered as three separate clades. Branch widths proportional to support (between 0.5 and 1.0).

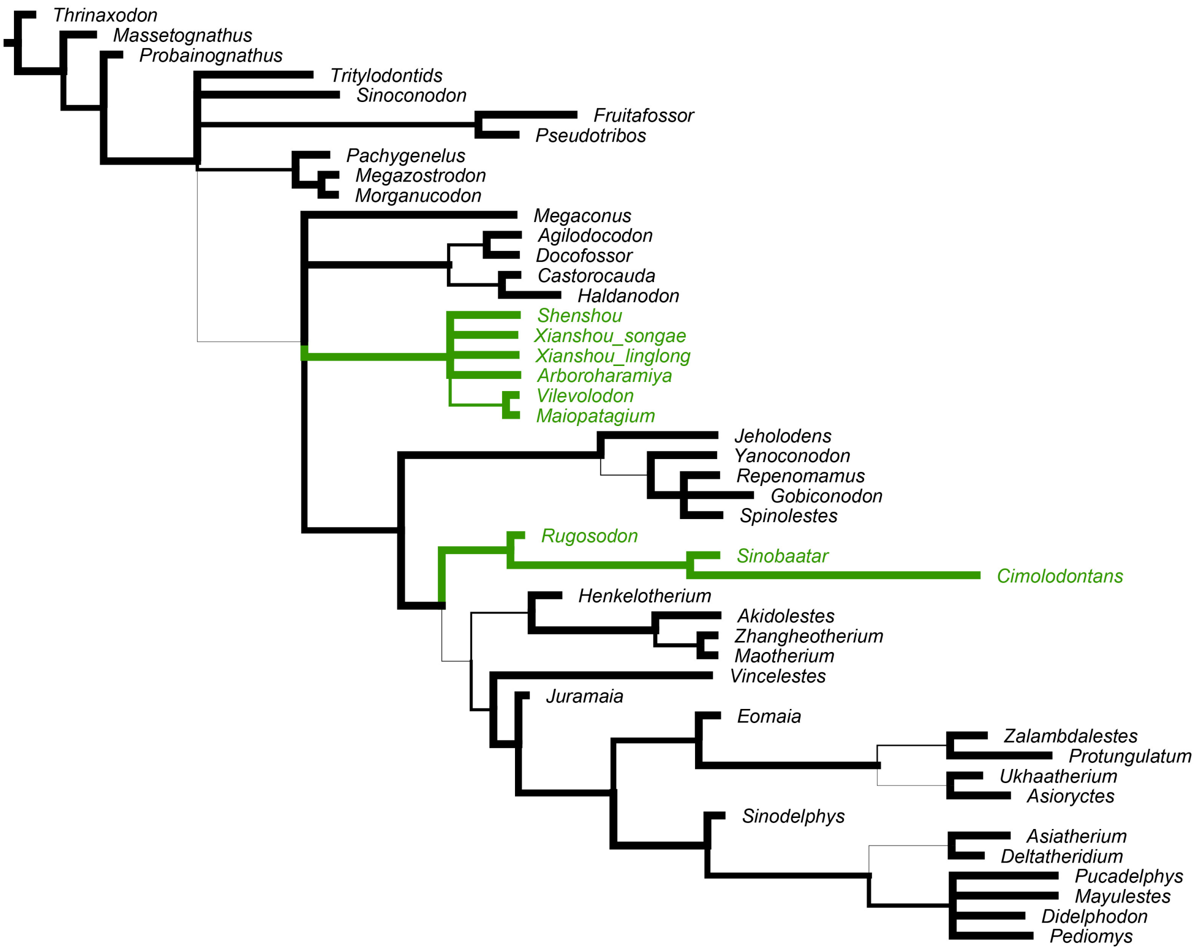


**Figure S5. Majority rule consensus tree of tip-dated analysis of postcranial characters from the dataset of Huttenlocker et al (2018).** This analysis recovers multituberculates separate from euharamiyids. However, relatively few taxa have postcranial characters preserved. Branch widths proportional to support (between 0.5 and 1.0).

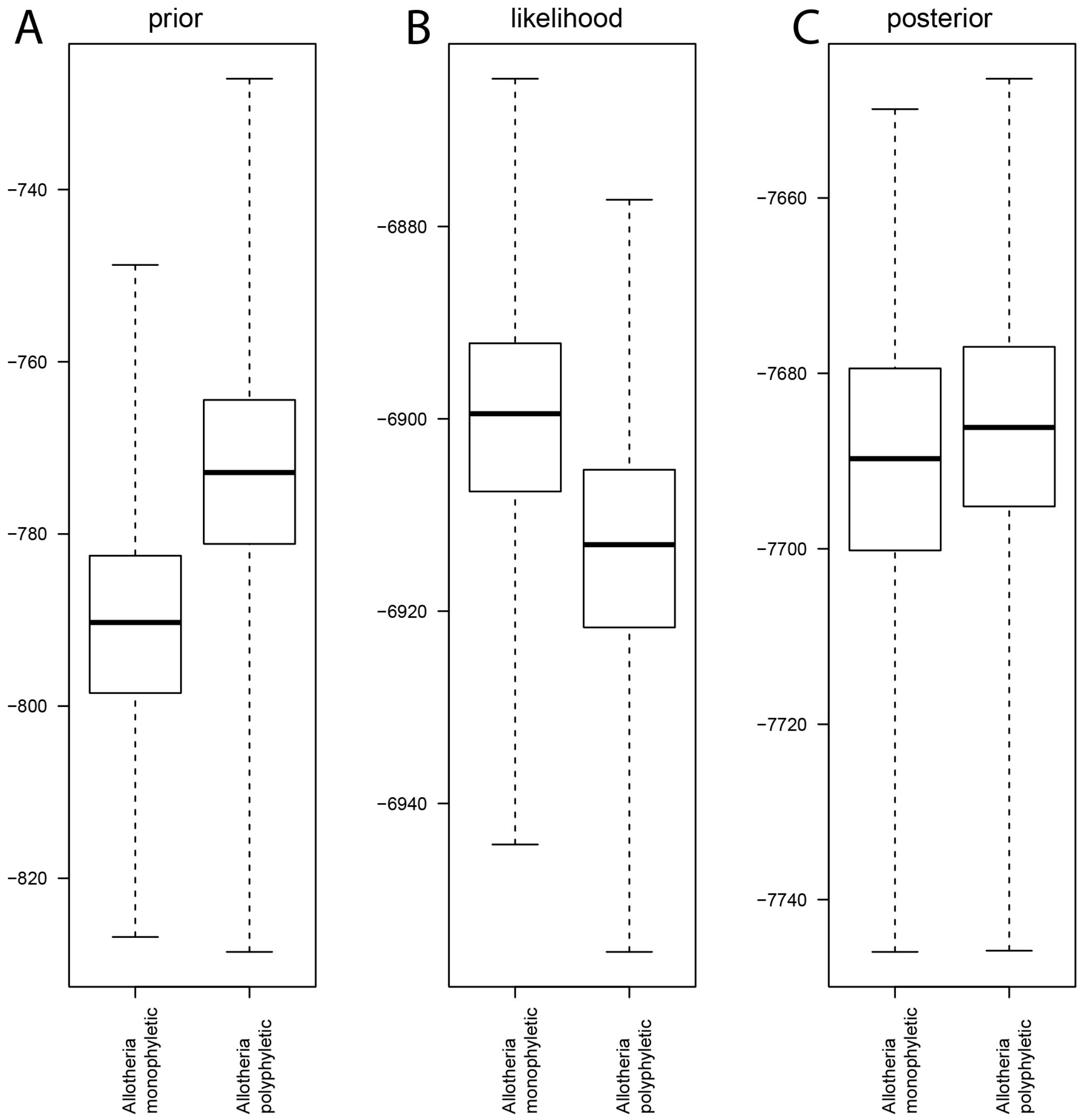


**Fig. S6. Analysis of the Krause et al. dataset shows twin peak behaviour associated with different phylogenetic positions for the haramiyids *Haramiyavia* and *Thomasia*.** A) Histogram of the prior probability in generations where Allotheria are sampled as monophyletic (left) or polyphyletic (right). The latter hypothesis generally has a higher probability. B) Histogram of the likelihood in generations where Allotheria are sampled as monophyletic (left) or polyphyletic (right). The former hypothesis generally has a higher likelihood. C) Histogram of the posterior probability in generations where Allotheria are sampled as monophyletic (left) or polyphyletic (right). The high prior – low likelihood of the latter and the low prior – high likelihood of the former almost cancel out, with only a slightly higher posterior probability in generations that sample allotherian polyphyly.

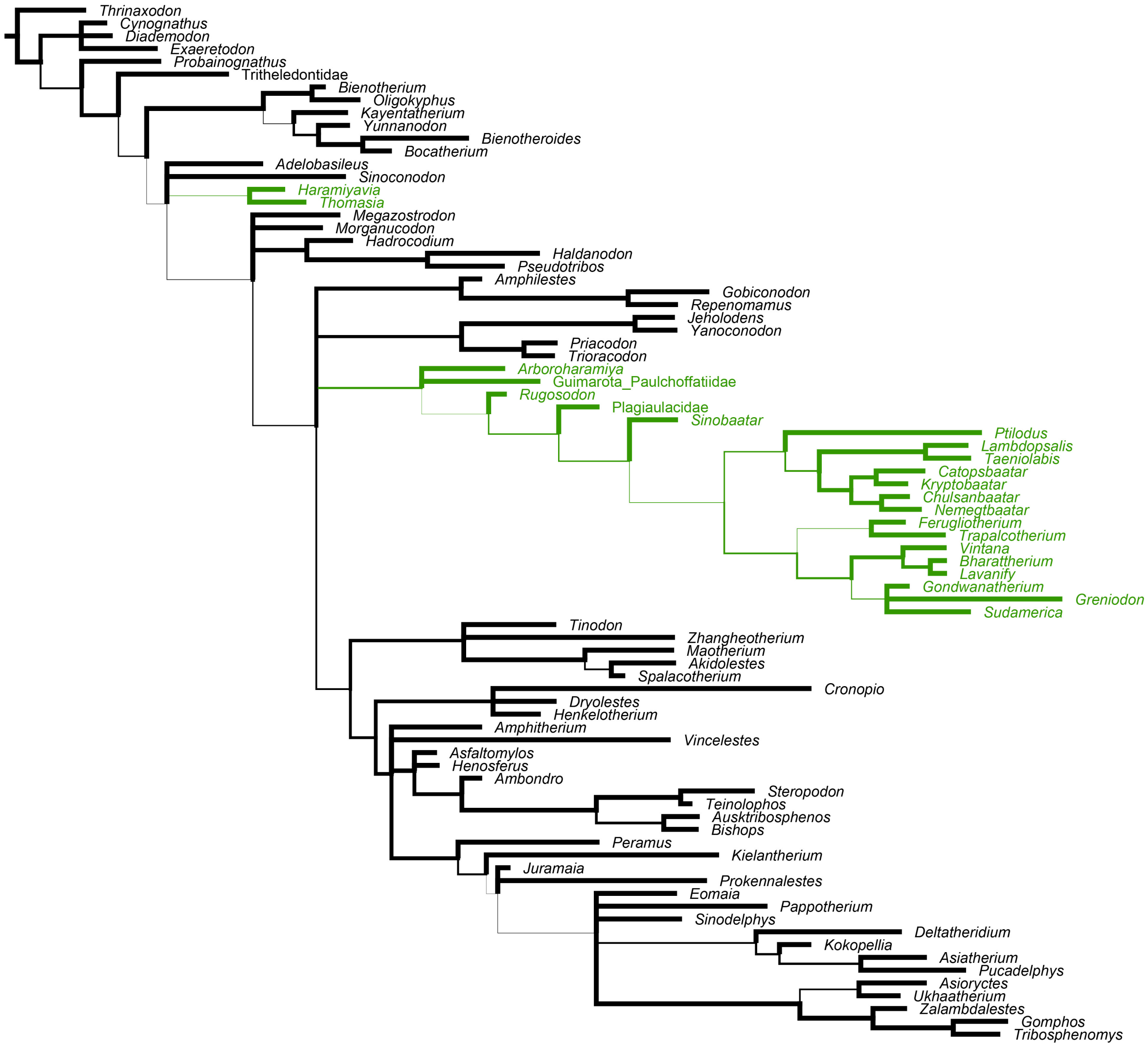


**Figure S7. Majority rule consensus tree from tip-dated analysis of the dataset of Krause et al. (2014).** Allotheria are recovered as polyphyletic with weak support. Branch widths proportional to support (between 0.5 and 1.0).

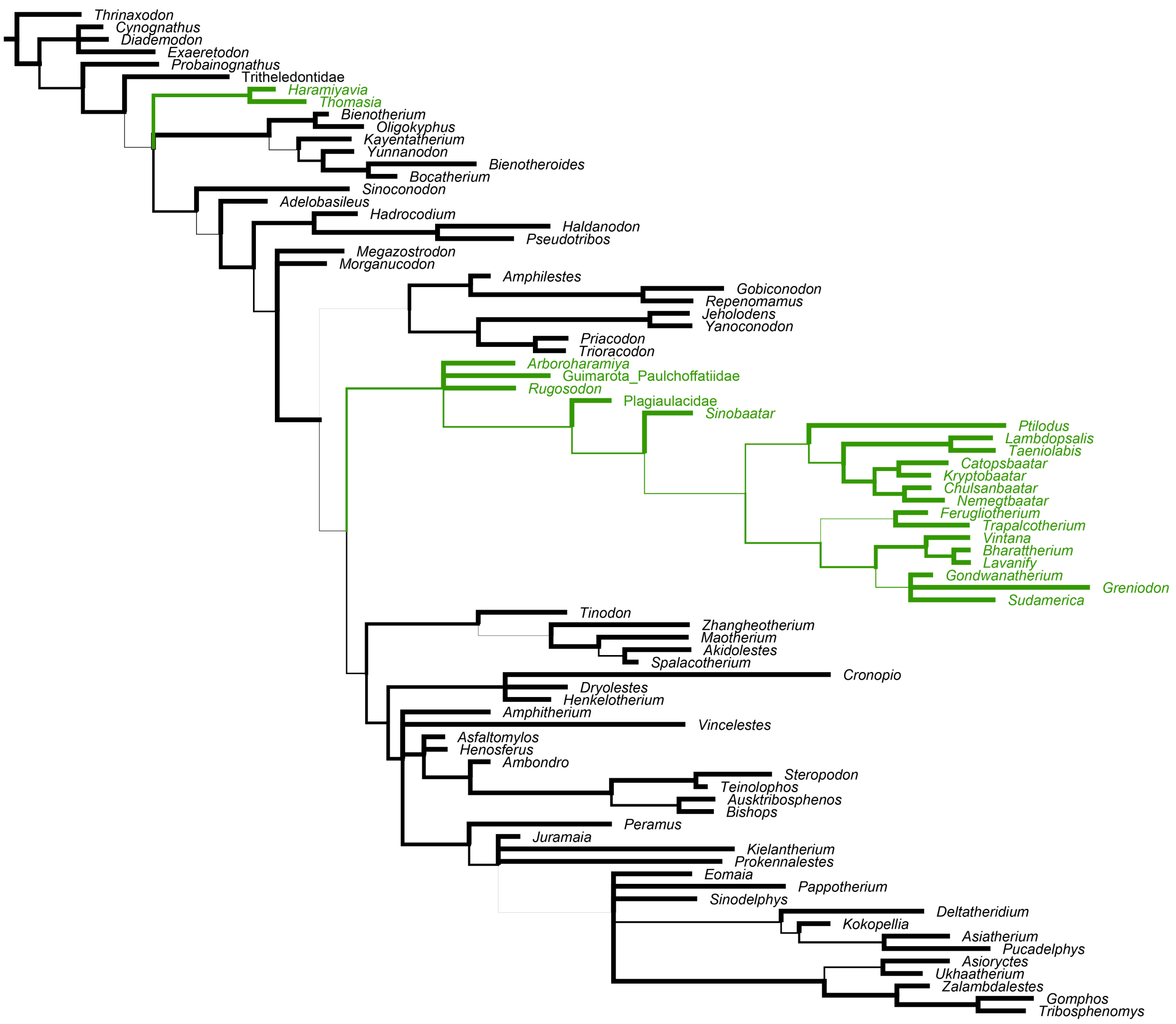


**Figure S8. Majority rule consensus tree from tip-dated analysis of the dataset of Krause et al. (2014), using only those generations in which allotheria were sampled as polyphyletic.** This consensus tree summarises 63% of the posterior sample. Branch widths proportional to support (between 0.5 and 1.0).

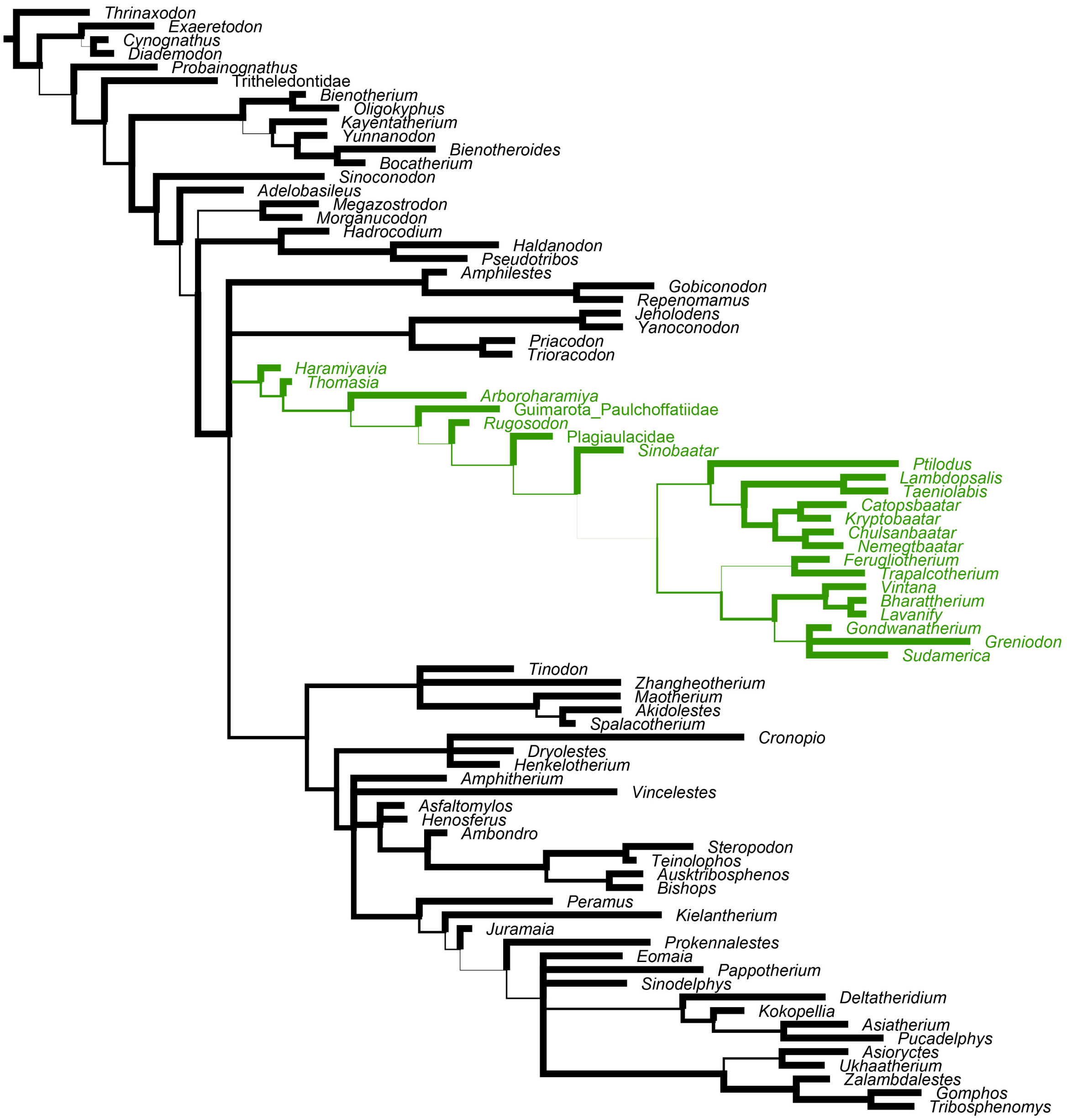


**Figure S9. Majority rule consensus tree from tip-dated analysis of the dataset of Krause et al. (2014), using only those generations in which allotheria were sampled as monophyletic.** This consensus tree summarises 37% of the posterior sample. Branch widths proportional to support (between 0.5 and 1.0). Weakly supported branches within Allotheria are due to the lability of the phylogenetic position of some or all gondwanatheres. Gondwanatheres were excluded from consideration for which generations of the Bayesian analysis sampled allotherian monophyly.

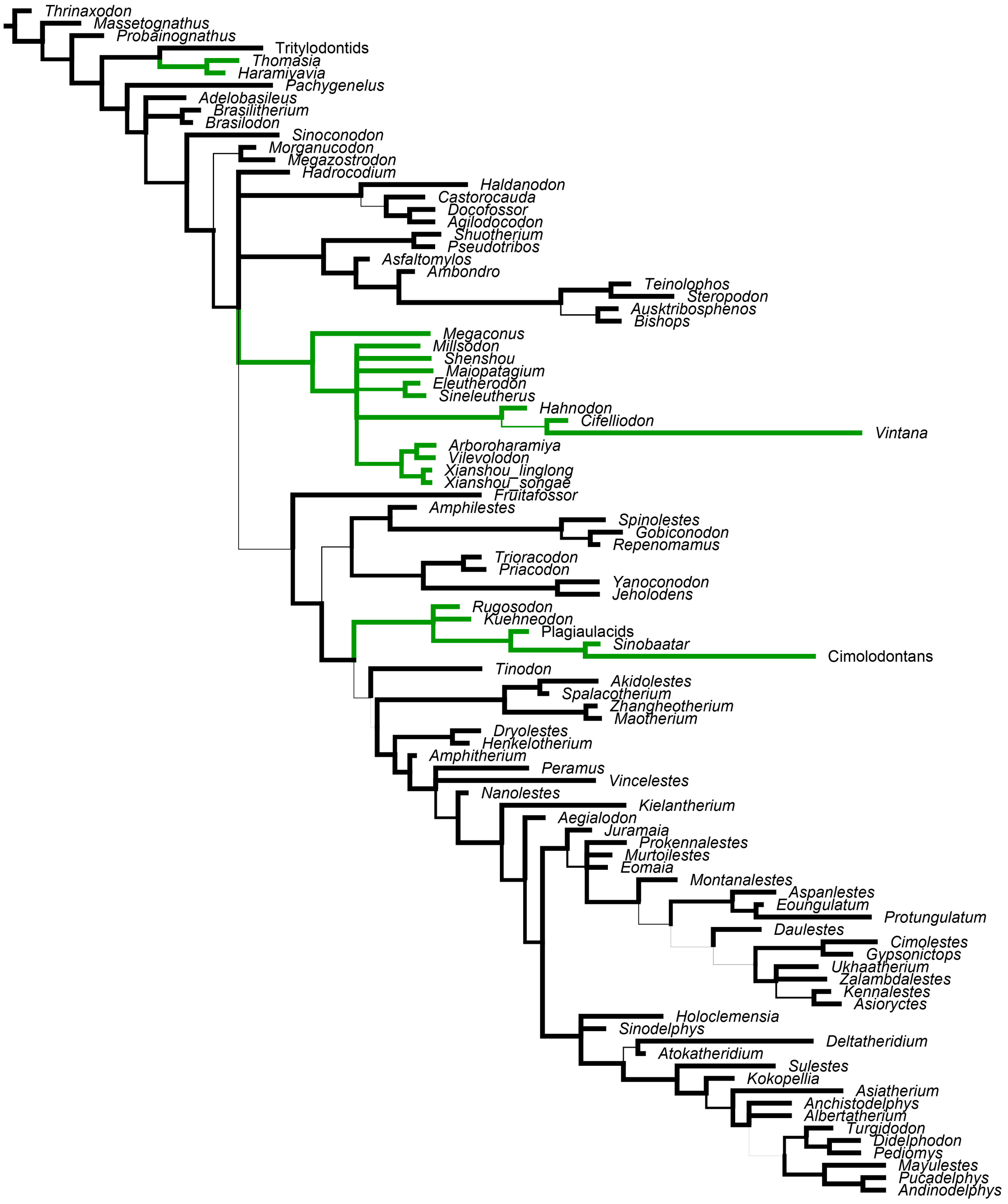


**Figure S10. Majority rule consensus tree from tip-dated analysis of the dataset of Huttenlocker et al. (2018), with a unrestrictive prior on the age of *Juramaia*.** The tree is congruent with the main analysis in which a Middle Jurassic age for *Juramaia* is used. Analysis with a less restrictive prior leads to strong support for a Lower Cretaceous age, but other aspects of the result are unaffected.

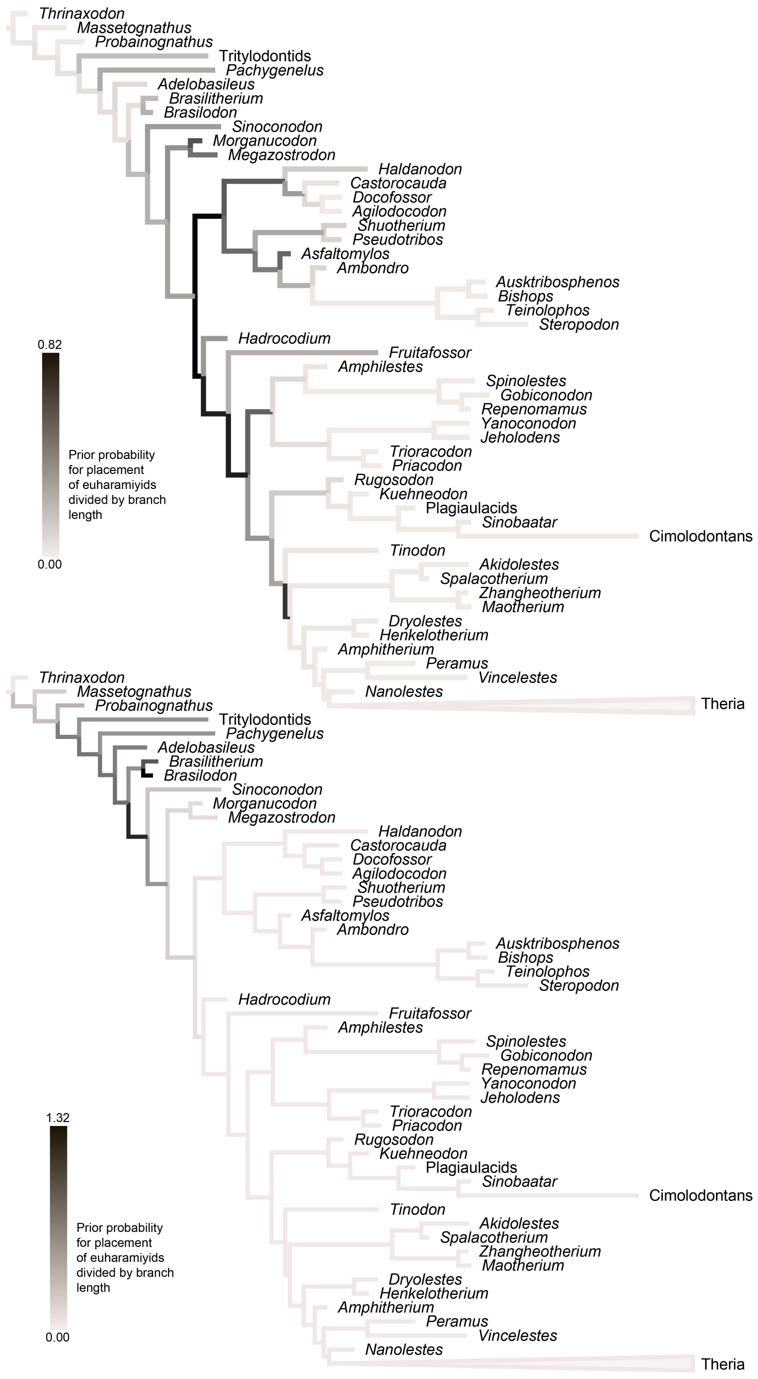


**Figure S11. Topology prior for haramiyids, corrected for branch length.** A-B) The tree is a fixed topology based on the maximum clade credibility tree from the main analysis, on which the prior probabilities for the position of two groups, divided by the branch length, of haramiyids are mapped. Branch colours represent the corrected prior probability that the respective clade (A, euharamiyids; B, Triassic haramiyids) was found on that branch in an analysis run without data. The corrected prior probabilities for Triassic haramyids are concentrated at the base of the tree, unlike the corrected prior for euharamiyids, which is more diffuse and centred in younger parts of the tree. Removing the effect of branch lengths clearly shows the temporal signal in the prior.

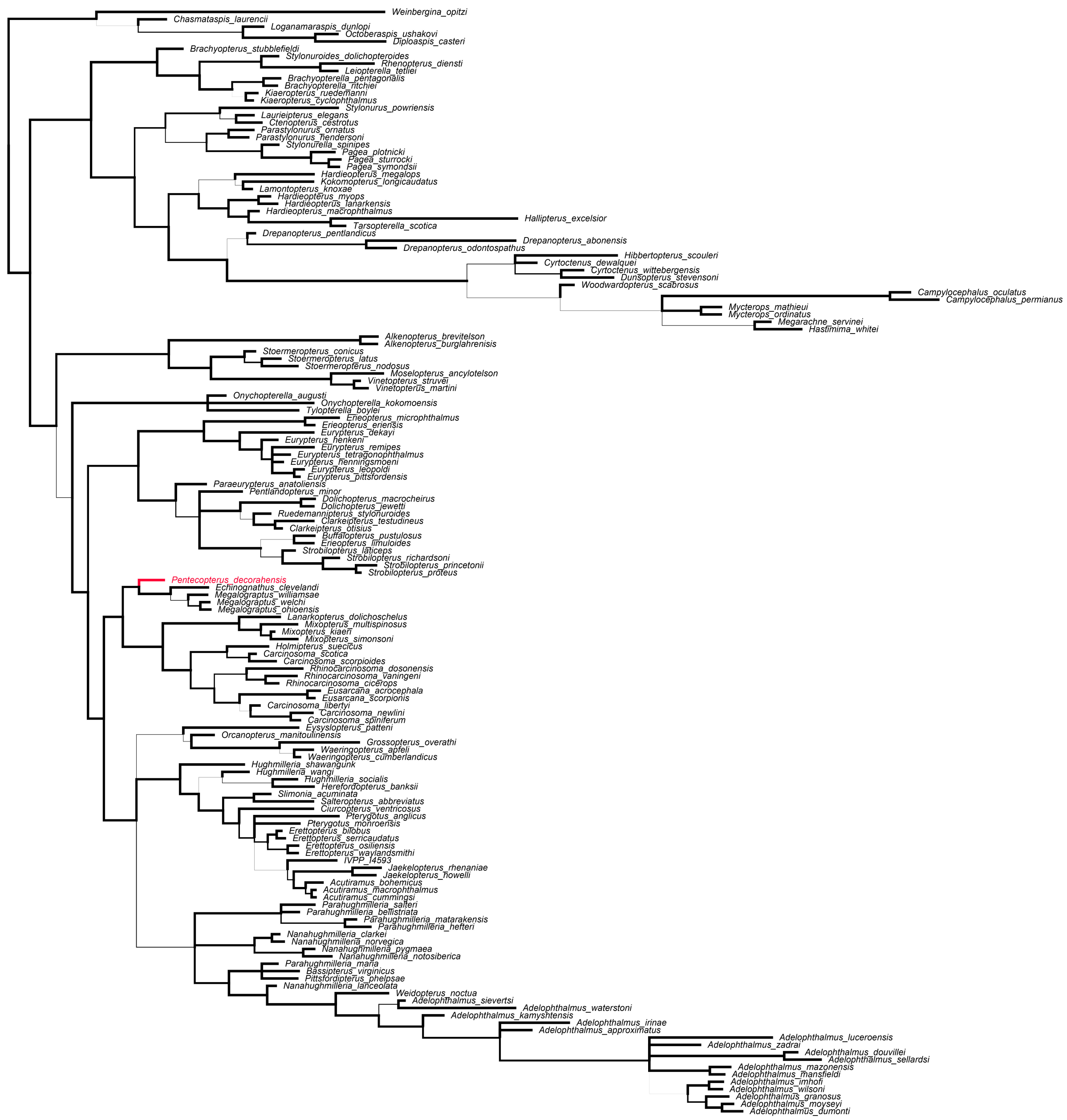


**Figure S12. Majority rule consensus tree from tip-dated analysis of the eurypterid dataset of Lamsdell and Selden (2016).** The oldest eurypterid, *Penecopterus*, is highlighted in red. This is found in a deeply nested, stratigraphically incongruent position. This shows that  tip-dating will not over-rule stratigraphically incongruent hypotheses supported by sufficient morphological data.
